# Prospective cohort study reveals unexpected aetiologies of livestock abortion in northern Tanzania

**DOI:** 10.1101/2021.08.23.457407

**Authors:** Kate M. Thomas, Tito Kibona, John R. Claxton, William A. de Glanville, Felix Lankester, Nelson Amani, Joram J. Buza, Ryan W. Carter, Gail E. Chapman, John A. Crump, Mark P. Dagleish, Jo E. B. Halliday, Clare M. Hamilton, Elisabeth A. Innes, Frank Katzer, Morag Livingstone, David Longbottom, Caroline Millins, Blandina T. Mmbaga, Victor Mosha, James Nyarobi, Obed M. Nyasebwa, George C. Russell, Paul N. Sanka, George Semango, Nick Wheelhouse, Brian J. Willett, Sarah Cleaveland, Kathryn J. Allan

**Affiliations:** Centre for International Health, Dunedin School of Medicine, University of Otago, Dunedin, New Zealand; Kilimanjaro Clinical Research Institute, Good Samaritan Foundation, Moshi, United Republic of Tanzania; Nelson Mandela African Institution of Science and Technology (NM-AIST), Tengeru, United Republic of Tanzania; Institute of Biodiversity, Animal Health & Comparative Medicine, College of Medical Veterinary and Life Sciences, University of Glasgow, Glasgow, United Kingdom; Paul G. Allen School for Global Animal Health, Washington State University, WA, United States of America; Global Animal Health Tanzania, Arusha, United Republic of Tanzania; School of Veterinary Medicine, University of Glasgow, Glasgow, United Kingdom; Kilimanjaro Christian Medical University College, Moshi, United Republic of Tanzania; Moredun Research Institute, Midlothian, United Kingdom; Zonal Veterinary Centre – Arusha, Ministry of Livestock and Fisheries, Arusha, United Republic of Tanzania; Tanzania Veterinary Laboratory Agency, Arusha, United Republic of Tanzania; School of Applied Sciences, Edinburgh Napier University, Edinburgh, United Kingdom; Medical Research Council, University of Glasgow Centre for Virus Research, Glasgow, United Kingdom

## Abstract

Livestock abortion is an important cause of productivity losses worldwide and many infectious causes of abortion are zoonotic pathogens that impact on human health. Little is known about the relative importance of infectious causes of livestock abortion in Africa, including in subsistence farming communities that are critically dependent on livestock for food, income, and wellbeing. We conducted a prospective cohort study of livestock abortion, supported by cross-sectional serosurveillance, to determine aetiologies of livestock abortions in livestock in Tanzania. This approach generated several important findings including detection of a Rift Valley fever virus outbreak in cattle; high prevalence of *C. burnetii* infection in livestock; and the first report of *Neospora caninum*, *Toxoplasma gondii,* and pestiviruses associated with livestock abortion in Tanzania. Our approach provides a model for abortion surveillance in resource-limited settings. Our findings add substantially to current knowledge in sub-Saharan Africa, providing important evidence from which to prioritise disease interventions.

## Introduction

Infectious causes of abortion, including early foetal loss and stillbirth, affect livestock production worldwide with the potential to cause major economic losses ^1^. Many abortions are caused by zoonotic pathogens that also pose a risk of infection to people ^2^. Although the economic losses can be substantial in high-production, intensive agricultural systems ^1^, abortion-associated livestock production losses have marked negative impacts for the poorest communities in the world, many millions of whom are critically dependent on livestock for economic livelihoods, food security, health, and wellbeing ^3, 4^.

Livestock abortions can be caused by a wide range of pathogens as well as non-infectious causes such as nutritional deficiencies and trauma ^2^. While many infectious causes of abortions occur worldwide, aetiological data based on robust laboratory-confirmed diagnoses are dominated by studies from farming systems in high- and middle-income countries ^5–7^ with data lacking from low-income settings.

Comparison of aetiologies across geographic areas is challenging as studies vary in methodologies and in the specific pathogens that are investigated. Furthermore, the ability to identify a specific aetiology of a livestock abortion is often relatively low, with less than 35% cases attributed to specific diagnoses ^5, 7, 8^.

Considerable surveillance and diagnostic infrastructure is required for robust case attribution of livestock abortions ^5, 9^, which typically limits the data available from low resource settings. For countries where data do exist, geographic patterns appear to vary quite widely. For example, *Coxiella burnetii* has been determined to be a common cause of abortion in sheep and goats in Canada ^10^, and in goats in the Netherlands ^11^, but only rarely in sheep in the United States of America (USA) ^8, 12^ or the United Kingdom (UK) ^9^, despite evidence of widespread seropositivity in both countries ^13, 14^. *Chlamydia abortus* is one of the most common causes of infectious ovine abortion worldwide, including the UK, the USA and most of northern Europe but has not been found to be an abortion issue in sheep in Australia or New Zealand ^15^ and there is little available information from the African continent, other than in a few select countries (e.g. Tunisia, Algeria, Namibia, Zimbabwe, and Morocco ^16–19^. In tropical areas, epidemic-prone pathogens, such as Rift Valley fever virus (RVFV) which poses a severe health threat to both animals and people, are also a concern ^20^. No less important are the endemic zoonoses such as *Brucella abortus* and *B. melitensis*, and zoonotic protozoal agents such as *Toxoplasma gondii,* which are of substantial concern in subsistence farming settings where people live closely with their animals ^3, 21^ or in communities where HIV predisposes the human population to severe infection e.g., with toxoplasmosis ^22^.

In Africa, inferences about the potential role and relative importance of abortigenic agents are often made from serological studies of livestock ^23–25^. This is due to a number of factors including a lack of national-level surveillance for syndromes such as abortions in livestock; restricted access to veterinary care for many subsistence farmers, who are therefore unable to investigate individual abortion events; and limitations in diagnostic laboratory capacity (e.g., little to no access to histopathology services or molecular diagnostic tests that could help to generate data on different aetiologies of abortion). While serological data is valuable to identify the presence of abortigenic agents circulating within a livestock population, there is also a need to recognise the limitations of such data and the inferences that can be made based on it ^26, 27^. Livestock seropositivity indicates only that an animal has been exposed to a pathogen and does not necessarily indicate current disease status or causality of an abortion event. Additionally, chronic low-level carriage of some abortigenic agents (e.g., *C. burnetii*) in healthy animals complicates attribution of abortion. To date, in sub-Saharan Africa, there remains an almost complete absence of published data on livestock abortion aetiologies determined through confirmatory diagnostic testing, resulting in a very weak evidence base on which to prioritise animal health interventions.

In Tanzania, livestock diseases are prioritised for prevention and control largely on the basis of the potential for transboundary spread and zoonotic impacts ^28^. A recent exercise to prioritise zoonotic diseases ranked two abortigenic agents, *Brucella* and RVFV amongst the top six zoonoses for prevention and control ^29^. While there is clear evidence that brucellosis and Rift Valley fever (RVF) are both diseases of public health concern in Tanzania ^30–32^, no data currently exist on the relative importance of these or other zoonotic pathogens, such as *C. burnetii*, *Leptospira* spp., *C. abortus* or *T. gondii*, as causes of livestock abortion. Nor are there any data available on the contribution of other abortigenic pathogens of global importance, including *N. caninum*, bluetongue virus and pestiviruses such as bovine viral diarrhoea virus (BVDV) and border disease virus (BDV). Without robust aetiological data, it is unlikely that current approaches to livestock disease prioritisation will reflect the true impact of abortigenic pathogens, with the risk that scarce resources for disease control and prevention will be targeted ineffectively and achieve sub-optimal productivity and public health outcomes.

We sought to address knowledge gaps around the aetiology of livestock abortion in Tanzania through a prospective cohort study conducted in three regions spanning a range of agro-ecological settings ^33^. We used a combination of field investigation and laboratory diagnosis of abortion events in cattle, goats, and sheep to determine aetiologies of livestock abortions for a targeted selection of known abortigenic agents. For these selected agents, we developed a set of case definitions to attribute abortion events to a specific aetiological agent based on molecular diagnostic assays in the absence of comprehensive histopathology data. In the same study area, we also carried out serological analyses of a large sample of livestock sera collected by cross-sectional studies in the same area to determine seroprevalence to the same range of abortigenic pathogens at a regional level.

## Results

Figure 1 shows the location of sampling for the serological cross-sectional and abortion cohort studies.

**Figure 1:**
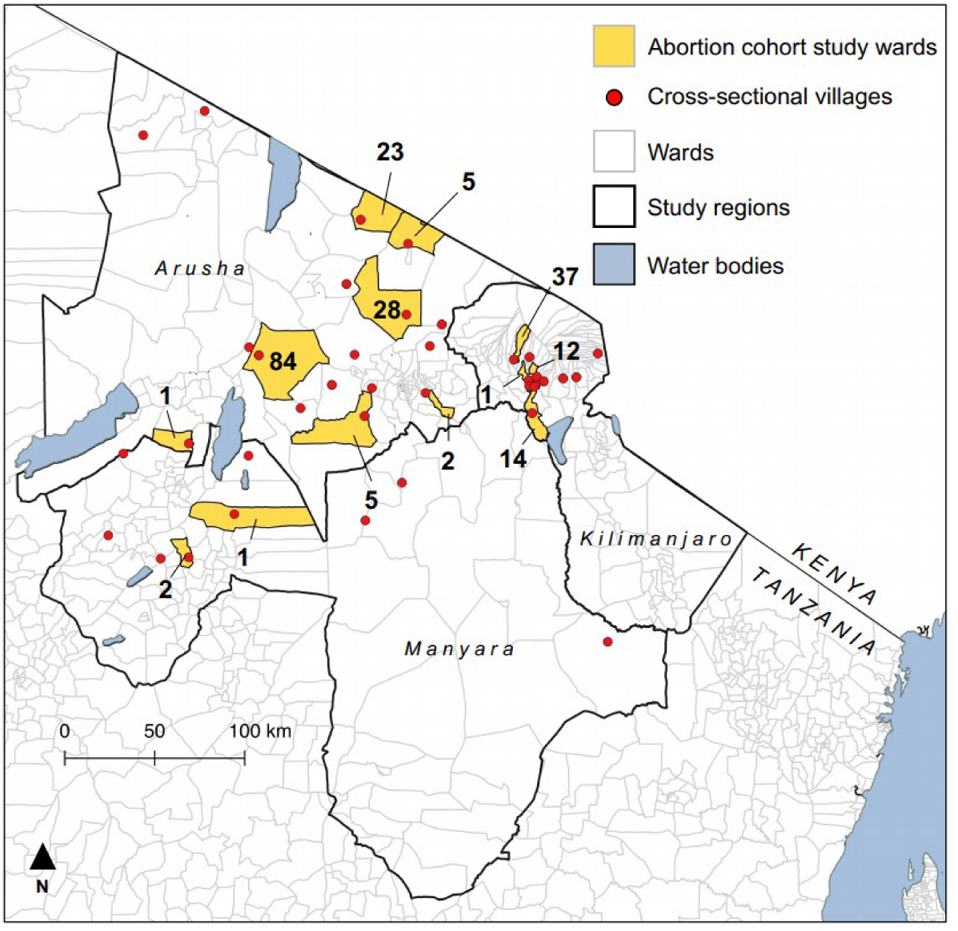
Distribution of villages in cross-sectional studies for serologic baseline testing (2013-2016; red dots) and abortion cohort sampling wards (2017-2019; yellow shading) with number of abortion events investigated, Arusha, Kilimanjaro, and Manyara Regions, northern Tanzania.

### Cross-sectional sample serology

Samples were available from 3,629 cattle, 3,344 goats, and 2,628 sheep from 567 households in 43 villages. Figure 2 summarises the cross-sectional seroprevalence of pathogens in ruminant livestock sampled in northern Tanzania. Seroprevalence at individual level for cross sectional livestock with Clopper Pearson 95% confidence intervals is detailed in supplementary materials (S1, A).

**Figure 2:**
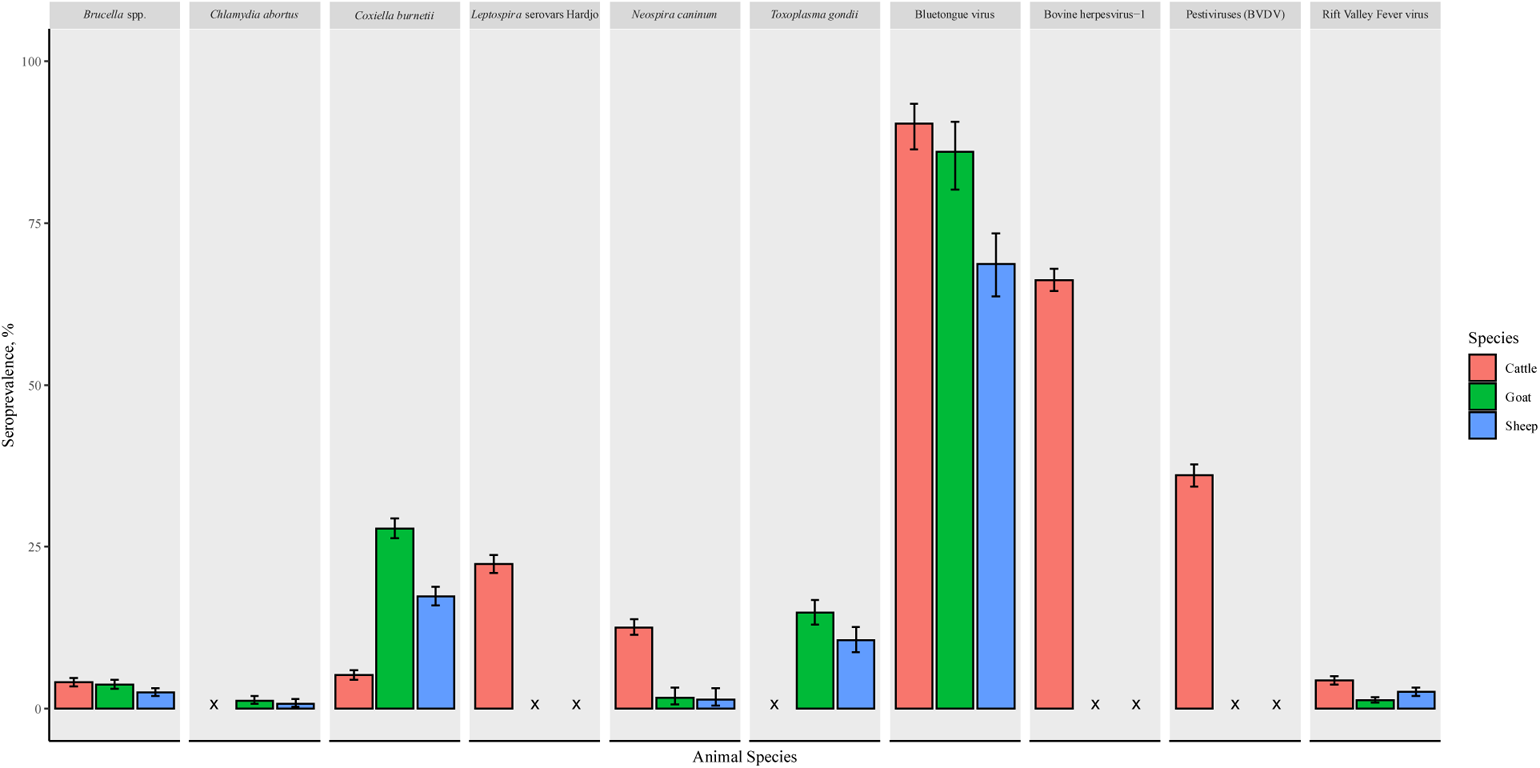
Seroprevalence of pathogens from cross-sectional studies (2013-2015 and 2016) in cattle, goats, and sheep, northern Tanzania. Where x = not tested and error bars represent Clopper-Pearson 95% confidence intervals

### Abortion cohort study

During the prospective cohort study period, a total of 215 abortion events were investigated, 71 (33%) in cattle, 100 (46.5%) in goats, and 44 (20.5%) in sheep. The numbers of each sample type collected for abortion events are detailed in Table 1.

**Table 1:** Number and type of samples collected from abortion events for each livestock animal species, northern Tanzania, 2017-2019

| Sample types collected | Cattle (n=71) | Goat (n=100) | Sheep (n=44) |
| --- | --- | --- | --- |
| Vaginal swabs | 71 (100%) | 98 (98.0%) | 44 (100%) |
| Exterior foetal swabs | 37 (52.1%) | 25 (25%) | 5 (11.4%) |
| Placental tissue | 34 (47.9%) | 25 (25%) | 1 (2.3%) |
| Acute sera | 71 (100%) | 98 (98.0%) | 43 (97.7%) |
| Convalescent sera | 49 (69.0%) | 56 (56.0%) | 29 (65.9%) |

### Abortion cohort acute serology

The acute seroprevalence results by pathogen in the abortion cohort was broadly similar to the cross-sectional survey. A table showing these results is available in the supplementary materials (S1; A).

### Diagnosis of abortion aetiology

We attributed 42 (19.5%) of 215 abortion events to one or more of the abortigenic pathogens included in our diagnostic panel. Table 2 summarises the number of abortion events with positive diagnostic test results that met the study case definitions. Histopathology and immunohistochemistry (IHC) results are shown in supplementary materials S1, B.

**Table 2:**
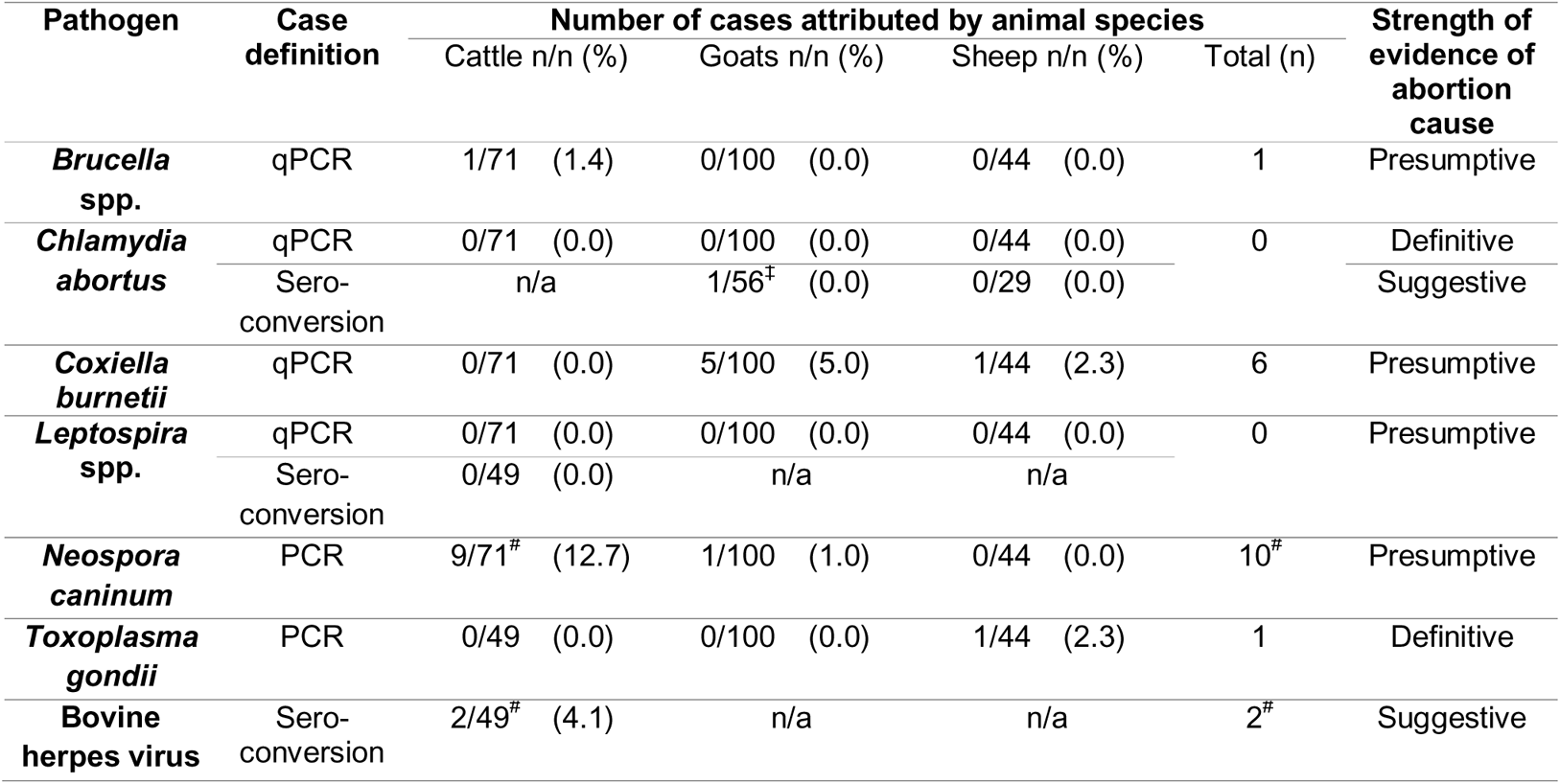
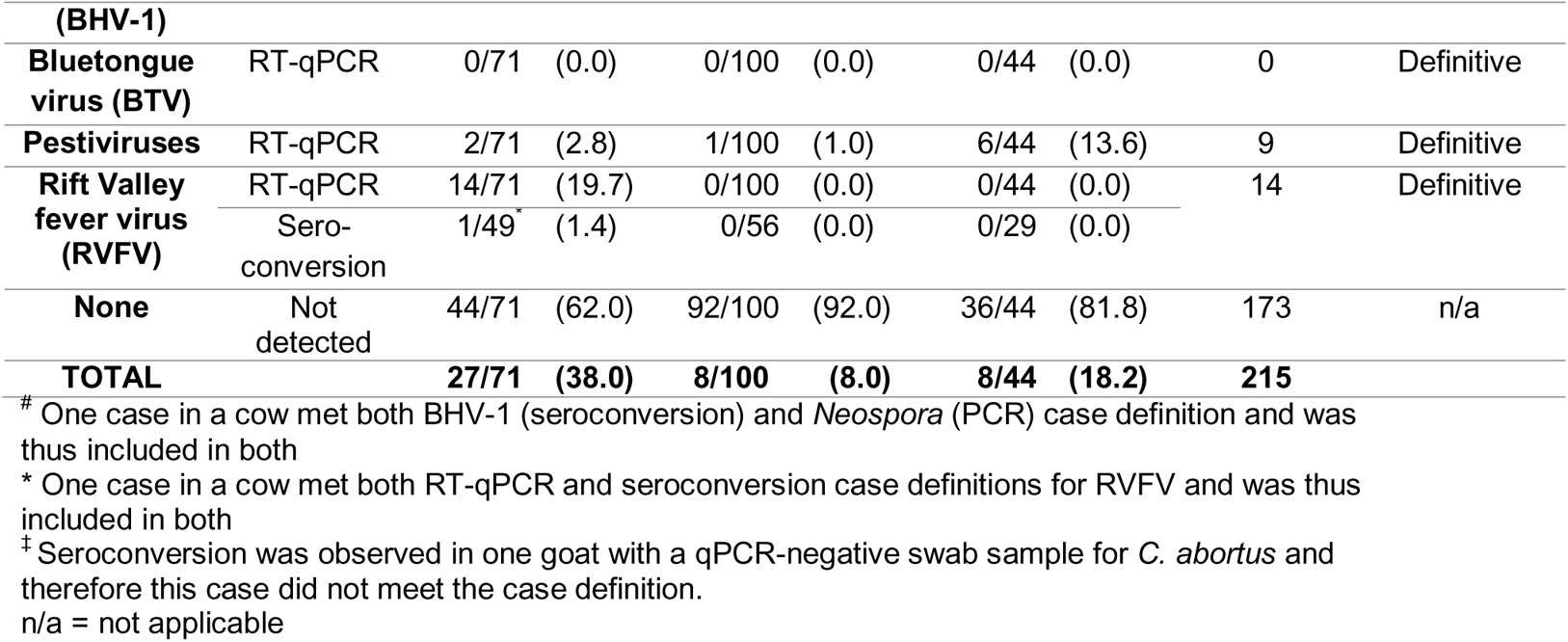
Number of abortion cases (cattle, goats and sheep) attributed to abortigenic agents using study case definitions, northern Tanzania 2017-2019

For some bacterial pathogens, despite evidence of infection in the dam, laboratory data was not considered sufficient to support a diagnosis of abortion associated with the pathogen. This was particularly true for *C. burnetii*, which was detected by qPCR with a threshold cycle (Ct) value <40 in 16 (22.5%) of 71 cattle, 24 (24.5%) of 98 goats, and 12 (27.3%) of 44 sheep, however only six (11.5%) of these 52 animals met the study case definition (Ct ≤ 27) for *C. burnetii* as a presumptive aetiology of abortion. For *C. abortus*, although seroconversion between acute and convalescent samples to *C. abortus* was detected in one (1.8%) of 56 goats with paired sera available for testing, this animal had a negative vaginal swab qPCR result and therefore did not meet our case definition for *C. abortus*-associated abortion.

Of the protozoal pathogens, *Neospora* DNA was diagnosed by nested conventional polymerase chain reaction (PCR) in nine (12.7%) of 71 cattle and one (1.0%) of 98 goat abortion events. In one cattle case, a co-infection with BHV-1 was also detected suggesting a multi-factorial aetiology. *Toxoplasma* DNA was detected by nested conventional PCR in one (2.3%) of 44 sheep abortion events. The single *T. gondii* PCR positive sheep sample was genotyped at 7/10 marker and was found to have type II alleles at all of them, indicative of a type II clonal genotype infection.

Of the viral pathogens, RVF was most commonly diagnosed, with RVFV RNA detected by reverse transcriptase qPCR (RT-qPCR) in 14 (19.7%) of 71 cattle abortion events. Evidence of seroconversion was detected in one animal that was positive by RT-qPCR. The other 13 RT-qPCR positive animals were positive in both acute and convalescent sera, meeting World Organisation for Animal Health (OIE) guidelines for definitive diagnosis of infection. Pestiviruses were also commonly detected with two (2.8%) of 71 cattle, six (13.6%) of 44 sheep, and one (1.0%) of 100 goat abortion events positive by RT-qPCR.

### Pestivirus speciation results

Among the nine animals that tested positive for pestiviruses by RT-qPCR, N^Pro^ coding sequence was obtained from six samples for phylogenetic analysis (Figure 3). BVDV infection was confirmed in two animals (one cow and one sheep) with a distinct genotype detected in each case. The cattle sample (SF1-190) was most similar to reference sequences from BVDV genotype 1d whereas the sheep sample (SD1-147) was most similar to reference sequences from BVDV genotype 1b.

**Figure 3:**
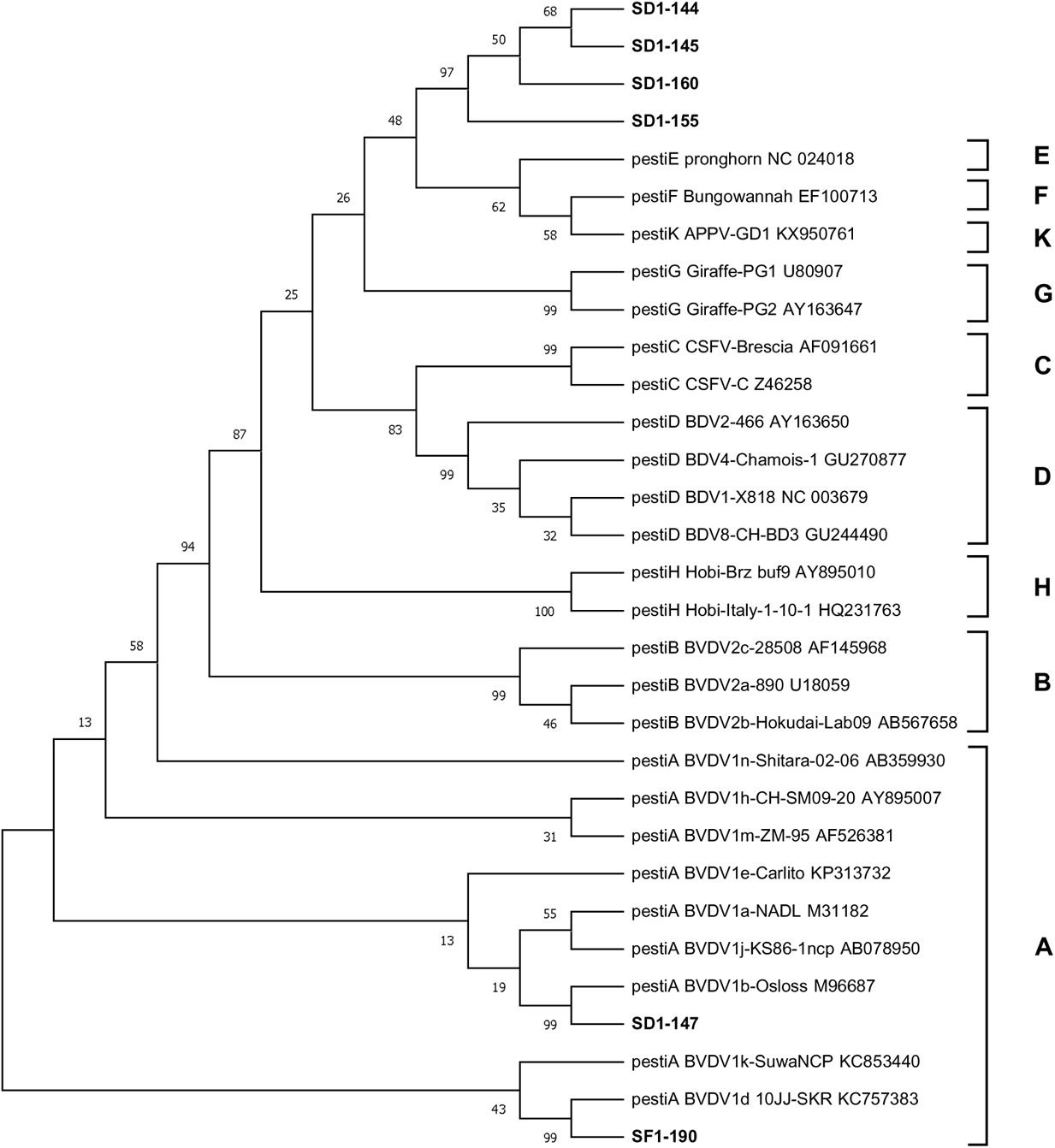
Results of evolutionary analysis of the N^Pro^ coding region from pestivirus sequences obtained from cattle and sheep sampled in the abortion cohort study, northern Tanzania.

Sequences obtained from the remaining four samples (SD1-144, SD1-145, SD1-155, SD1-160), all obtained from sheep, were similar to each other but appeared to be distinct from all of the reference sequences. These sequences represent a potentially novel, previously unreported pestivirus species and formed a separate clade on phylogenetic analysis. The sequences were assigned accession numbers OU548650-OU548655 by the European Nucleotide Archive (https://www.ebi.ac.uk/ena/browser/home) and, for the purposes of that submission, this novel clade was named Pestivirus abortion/sheep/SD1/2018/Tanzania.

The evolutionary history was inferred using the Maximum Likelihood method and Tamura-Nei model of nucleotide substitution ^34^. The tree with the highest log likelihood is shown. The percentage of trees from 500 Bootstrap repeats in which the associated taxa clustered together is shown next to the branches. Evolutionary analyses were conducted in MEGA X ^35^. Reference sequences representing nine pestivirus species were also included ^36^ and are labelled with virus species (A-H, K), genotype and strain name and Genbank accession numbers. Pestivirus species are also shown by bracket labels on the right of the figure. Samples analysed in this study are shown in bold and labelled with unique identifiers (SD/F1-XXX), where SD denotes vaginal swab taken from aborting dam; SF denotes swab taken from aborted foetus. Abbreviations: APPV = atypical porcine pestivirus; BDV = border disease virus; BVDV = bovine viral diarrhoea virus; CSFV = classical swine fever virus.

## Discussion

To our knowledge, this is the first multi-pathogen study of the aetiologies of livestock abortion in Tanzania. We developed and applied an approach designed to optimise abortion investigations in resource limited settings, complemented by traditional cross-sectional surveillance to generate a more comprehensive understanding of the distribution and impact of several important abortigenic pathogens. This approach generated several new insights for Tanzania including: a) detection of an RVF outbreak in cattle during an inter-epidemic period; b) relatively high prevalence of *C. burnetii* infection in cattle, goats and sheep; c) the first demonstration of *N. caninum*, *T. gondii,* and pestiviruses, including novel pestivirus species, associated with livestock abortion in Tanzania, including detection in less typical hosts (for example, *N. caninum* in goats); but, contrary to prior expectations, d) limited or no detection of *Brucella* spp., *C. abortus*, and *Leptospira* spp. in cases of livestock abortion. Our findings provide important information on the health and production impacts of a range of livestock pathogens in East Africa with relevance for prioritising disease interventions in the region.

Our study demonstrates the importance of zoonotic pathogens as causes of abortion in ruminants in Tanzania, with implications for human as well as animal health. Two important zoonotic pathogens were diagnosed in multiple abortion events by our study: RVFV, a virus that can cause devastating outbreaks of human and animal disease ^37, 38^, and *C. burnetii*, an important bacterial cause of human febrile illness in Tanzania ^39^. RVFV is a high priority zoonosis in East Africa and outbreaks are frequently associated with livestock abortion storms ^20^. Direct animal to human transmission of infection can occur following inhalation of aerosolized virus particles or during close contact with tissues and bodily fluids of infected animals, for example, when assisting with parturition or handling aborted foetuses ^40, 41^. Although RVF usually occurs as large outbreaks occurring every 10-20 years, there is increasing serological evidence of transmission in the period between these epidemics ^42^. This study detected the occurrence of an apparent outbreak in cattle that was previously unreported in Tanzania, despite RVF outbreaks being detected in Kenya, Uganda, and Rwanda at the same time ^40, 43^. Although an alert of RVF was issued in Tanzania in June 2018 at the time of this study ^44^, to our knowledge, no human cases were reported. Our finding highlights the value of livestock abortion surveillance in early detection of this epidemic-prone pathogen and also demonstrates the value of molecular diagnostic capacity for detection of RVF livestock cases.

*Coxiella burnetii* is a pathogen of concern in Europe, where outbreaks of infection in small ruminant livestock have also led to outbreaks of human disease ^45^. *C. burnetii* has also been detected in humans and in a wide range of animal species across Africa ^46, 47^, and accounts for notable proportions of severe human febrile illness in Africa (Vanderburg et al., 2014), including in northern Tanzania ^39, 48^. Our results provide important evidence of the impact of *C. burnetii* as a cause of livestock reproductive losses, particularly in small ruminants, revealing a dual human and animal health burden of this zoonotic pathogen. In our abortion cohort study, we applied a stringent case definition that exceeded the minimum guidelines stated by the OIE for diagnosis of *C. burnetii-*associated abortion. Using this case definition, we diagnosed six cases of *C. burnetii-*associated abortion, but it is also worth noting that *C. burnetii* DNA was detected in a total of 52 (24.4%) abortion events indicating that shedding of this zoonotic bacterium is even more prevalent than the number of attributed cases would suggest. Based on our findings, we suggest that the development of preventive and control measures against *C. burnetii* should be a priority for future work that could have substantial benefits for human and animal health.

Our study confirmed *T. gondii,* another zoonotic pathogen, is associated with livestock abortion in Tanzania. Toxoplasmosis is also an important, sometimes fatal, zoonosis in Tanzania, particularly among immunocompromised people ^22^. There are limited data on genotypes present in Africa but the Type II clonal lineage found in the single PCR positive sheep abortion event is known to be dominant in North and East Africa so aligns with previous studies ^49^. The cross-sectional serological data showing widespread exposure of *T. gondii* in sheep and goats are consistent with previous published findings of high seroprevalence in people and animals, including livestock ^50, 51^ and wild felid species in East Africa ^52^. Given that toxoplasmosis is one of the most important abortigenic agents of sheep worldwide, it is perhaps surprising that we did not detect more cases of *T. gondii* as a cause of sheep abortion in this study. However, high flock seroprevalences are typically associated with low rates of abortion due to the development of protective immunity in ewes infected with *T. gondii* prior to mating ^53^.

Other zoonotic pathogens were detected infrequently in the abortion cohort study, with only a single case attributed to *Brucella* spp. in a cow, a single detection of seroconversion to *C. abortus* in a goat, and no evidence of abortions associated with *Leptospira* spp. in any livestock species. Although *C. abortus* has been reported as a cause of abortion in sheep elsewhere in northern and southern Africa ^54–56^, our study generated little evidence that *C. abortus* was contributing to the burden of production losses in sheep in northern Tanzania at the time of our study. Our cross-sectional serological findings also support the conclusion that levels of exposure in small ruminants are very low across northern Tanzania. However, these findings do raise concerns about the potential impact should infection be introduced to this predominantly naïve population in the future.

Both *Brucella* spp. and *Leptospira* spp. are recognised as important zoonoses in northern Tanzania, causing large proportions of severe human febrile illness ^21, 48, 57, 58^ and are known to be circulating in livestock in these areas of Tanzania. Seroprevalence of *Brucella* spp. was low in all three livestock species (Fig. 2), which is consistent with the low frequency of detection in abortion cases. In contrast, although *Leptospira* was not detected by qPCR in any abortion event investigated, evidence of exposure to *Leptospira borgpetersenii* serovar Hardjo, and *Leptospira interrogans* serovar Hardjo (subsequently referred to as *Leptospira* Hardjo serovars) was relatively high in cattle in the cross-sectional study. Several possible explanations may explain this discrepancy. Similar to *T. gondii,* prior exposure to *Leptospira* serovars may lead to protective immunity prior to animals becoming pregnant. Additionally, livestock abortions associated with *Leptospira* are often seen in outbreaks or ‘abortion storms’ following either a new introduction into a herd or a seasonal outbreak where flooding or wet weather conditions support transmission of novel serovars ^59^. The timespan of our project meant that we may have missed intermittent or infrequent abortion storms associated with *Leptospira* in livestock.

However, even accounting for these limitations, our study suggests that in these agricultural settings, the proportion of abortions attributable to *Brucella* spp. and *Leptospira* spp. may be lower than is widely perceived. Alongside other constraints to the uptake and use of livestock brucellosis vaccines these data raise questions about the likely effectiveness and sustainability of livestock vaccination-based control strategies in this context if based on a farmer pays model only. To effectively mitigate both the animal and human impacts of brucellosis a diversity of strategies may be needed to achieve the vaccination coverage required to reduce animal production impacts and also to reduce the substantial human health burden from this disease.

In addition to zoonotic pathogens, our study provides also provides valuable data on several production-limiting infections of ruminants that are often over-looked in sub-Saharan Africa. In particular, while *N. caninum* and pestiviruses are considered important causes of abortions and productivity loss in Europe and the USA, there are few data available describing their impact in African livestock. Our results provide the first direct evidence of livestock abortion associated with *N. caninum* infection in Tanzania, a conclusion which is supported by a positive relationship between *N caninum* seroprevalence and in-herd abortion rates in cattle detected by earlier work in our study area ^60^. In this study, *N. caninum* was predominately diagnosed among cattle abortions, but notably also in one goat abortion event. Serological evidence of low-level exposure to *N. caninum* in goats (1.7%) and sheep (1.5%) was also detected across the region. Although abortion and foetal pathology associated with *N. caninum* has been described in goats and sheep ^61–63^, natural infection of *N. caninum* in small ruminants is not widely reported. Given the importance of small ruminants within livestock economies and for food security in Africa ^64, 65^, further investigation of the patterns and impact of *N. caninum* infection in small ruminants are warranted.

Pestiviruses were the most commonly diagnosed pathogen associated with abortion in sheep in this study, but pestiviruses are rarely discussed as priority production-limiting pathogens in livestock in Africa. Analysis of the six pestivirus sequences obtained in this study identified at least three different viral lineages associated with abortion in livestock in northern Tanzania including a novel lineage in sheep. Based on our analysis of the N^Pro^ coding region, this novel lineage appears to be distinct from other pestiviruses and is likely to be a new, previously unreported pestivirus species. Notably, several of the most similar pestivirus species were detected in sympatric wildlife species including giraffe in Kenya (Genbank accession numbers U80907 and AY163647). As wildlife interactions with livestock are common in many areas of Tanzania, we hypothesise that spill-over from wildlife may also be involved in the epidemiology of this novel pestivirus lineage, which appears to be a contributing cause of ovine reproductive loss in northern Tanzania.

Serological data from the cross-sectional study confirmed widespread exposure to BHV-1 in the study area (Fig. 2), and previous studies in this area have demonstrated an association between seropositivity and cattle abortion ^66^. However, the evidence for BHV-1 as a cause of abortion in this study is relatively weak. As latent BHV-1 infection may lead to false positive results by molecular detection ^67^, we relied on serology to diagnose infection. Two cattle showed evidence of BHV-1 seroconversion suggesting recent infection which may be implicated in the aetiology of abortion in these cases but further data are needed to evaluate the impact of BHV-1 pathogen on livestock reproduction in Tanzania and our findings highlight the challenges of attribution in settings where endemic pathogens may co-exist in cattle herds ^68^.

BTV, which causes production losses in livestock in Europe ^69, 70^, was not detected by RT-qPCR in any samples from livestock abortion events, despite high BTV seroprevalences detected in all species in the cross-sectional study population. Serological evidence of infection is widespread East Africa ^71–73^ and several studies have also demonstrated viraemia using RT-qPCR assays, reporting infection prevalence as high as 88.9% in cattle in Kenya ^73^ and 56.0% in goats in Uganda ^72^. Collectively, these studies suggest that BTV infection is endemic in livestock in East Africa. However, research indicates that African breeds of livestock may be relatively resistant to clinical disease associated with BTV. Studies from southern Africa have indicated that indigenous African breeds of livestock are less susceptible to clinical disease from BTV infection than European breeds ^74^. Furthermore, transplacental foetal infection is only associated with certain BTV strains, especially attenuated vaccine strains ^75^. Our findings supported by these trends in BTV epidemiology and pathogenesis suggest that BTV is not a major cause of livestock abortions in northern Tanzania and underscores the need for caution in making inferences about livestock diseases and causes of abortion in particular based solely on seroprevalence data.

The approach described in our abortion cohort study enabled us to attribute approximately 20% of reported abortion events to a specific aetiology. Our success rate was not dissimilar to the quoted rate of aetiological abortion diagnosis in high income settings (35%) ^5, 7, 8^ despite working in a challenging and resource-limited environment. Our approach was specifically designed to optimise abortion investigations in a resource-limited setting, where veterinary surveillance may be limited. Our focus on the use of molecular diagnostics to detect a range of selected pathogens proved to be valuable, particularly for high priority pathogens such as RVFV. While molecular diagnostic capability was established for several known abortigenic agents, it was not possible for our diagnostic panel to be fully comprehensive of all possible aetiologies of abortion and did not include agents such as *Campylobacter spp*., *Listeria spp*., and *Salmonella enterica* serovars that are known to be important causes of livestock abortion worldwide. Non-infectious causes were also not considered for investigation and these two factors may account for some of the undiagnosed cases.

Although histopathology is conventionally considered the most robust approach for attributing causality in livestock abortion, histopathology was of limited value in this study due to the limited availability and poor quality of tissues samples obtained through field investigations that were often conducted in remote settings. Additional challenges to inference of causality were encountered due to the nature of our study site. In areas such as rural Tanzania, where remote locations, high ambient temperatures and scavenging animals mean that the type and quality of diagnostic samples is often sub-optimal, inference about the cause can be particularly challenging. For example, placental tissues were not available for sampling from many small ruminants, and even where placental tissues were available for collection, most tissue samples showed evidence of significant autolysis, which precluded a definitive diagnosis in some cases. However, despite these sampling limitations, molecular diagnostics provided a wealth of information regarding the presence of abortigenic agents in swab and tissue samples, and therefore we recommend that molecular diagnostics should be included as a core element of abortion surveillance studies to infer aetiology in resource-limited settings.

In considering appropriate criteria for attributing the aetiology of abortion events reported to this study, we became aware of variations in case definitions and the limitations of the evidence base underpinning attribution criteria for livestock abortions. While methods for assigning attribution in situations of imperfect diagnostic testing are well-developed and widely adopted in aetiological studies of human diseases ^76^, these questions are rarely addressed in veterinary investigations. Attribution was particularly problematic for *C. burnetii*, as the pathogen is frequently detected in vaginal fluids at parturition in clinically healthy animals ^77^ and environmental contamination of placental material or foetal surfaces may also lead to false positives in high risk environments ^78^. In this study, we developed a case definition for *C. burnetii* based on OIE guidelines ^79, 80^ but also driven by population-level data generated by this study. Further development of this case definition would be beneficial to understand how clinical history, gross pathology (where available), measured bacterial load and other adjunct diagnostics could be used to more confidently attribute abortion cases both within and outside the Tanzanian context.

Substantial differences in reporting rates between wards further limits the representativeness of our sample. Important differences in livestock seroprevalence between production systems have been observed in northern Tanzania ^81^ as well as in other areas of East Africa ^82^. Spatial heterogeneity in livestock exposure risk, with clustering in seropositivity at the village-level, have also been described for pathogens such as *Brucella* ^83^. It is therefore possible that sample collection in more administrative areas, with improved representation of pastoral and agropastoral livestock keepers, would reveal different frequencies of pathogen detection than described here. Our data should therefore be considered as only indicative of the relative importance of different pathogens as causes of livestock abortion in northern Tanzania as a whole.

A key insight from our study is that, apart from RVF, none of the abortigenic pathogens identified by this study have been considered important pathogens in Tanzania, or prioritised for interventions. Gaps remain in our understanding of the causes of livestock abortion in Tanzania. However, even with testing a limited range of pathogens, our study has been able to generate data that adds substantially to the existing evidence base for the development of interventions to reduce the burden of production losses associated with livestock abortions. Our study further highlights the value and importance of establishing laboratory-based molecular diagnostics and developing clear case definitions and diagnostic criteria for use in resource-limited settings. With laboratory capacity and technical capability expanding rapidly in Africa, there is enormous potential and an urgent need for the veterinary sector to shift towards more evidence-based policy and decision-making, with the potential for this transition to lead to substantial improvements in animal and human health.

## Methods

### Research ethics

Ethics approval for this research was granted by Kilimanjaro Christian Medical Centre (KCMC) Ethics Committee (No. 535 & No. 832); National Institute of Medical Research (NIMR), Tanzania (NIMR/HQ/R.8a/Vol.IX/1522 & NIMR/HQ/R.8a/Vol.IX/2028); Research Ethics Coordinating Committee and the Institutional Review Board for Clinical Investigations of Duke University Health System in the United States (Pro00037356), University of Otago Ethics Committee (H15/069 & H17/069); and University of Glasgow College of Medical Veterinary and Life Sciences Ethics Committee (200140152 & 200170006). Sample shipments between the UK and Tanzania were performed in accordance with the Nagoya Protocol on Access to Genetic Resources under Material Transfer Agreements between KCMC and University of Glasgow, and between KCMC and Moredun Research Institute (MRI) with authorisation for export and import, respectively, from the Tanzanian Ministry of Livestock and Fisheries (permit number VIC/ZIS/8582), and The Scottish Government’s Agriculture, Food and Rural Communities Directorate (TARP(S) 2019/07).

### Cross-sectional serological study

Two cross-sectional livestock serological studies were performed from 2013 through 2016, to provide a baseline of exposure to a range of abortigenic pathogens. Livestock were selected and sampled as previously described ^60, 84^ from six districts in Arusha Region, three districts in Kilimanjaro Region, and four districts in Manyara Region. Serum was separated from coagulated whole blood samples by centrifugation (≤1,300 x g for 10 mins at room temperature) and aliquoted to create replicate sets. Samples were transported to the Zoonoses Laboratory at Kilimanjaro Clinical Research Institute (KCRI), Moshi, Tanzania, for archiving at −80°C prior to testing. Cattle, goat, and sheep sera were tested for exposure to *Brucella* spp., *C. burnetii,* and RVFV using commercial ELISA, as per manufacturer’s instructions (Table 3). Cattle sera were additionally tested for exposure to *Leptospira* Hardjo serovars, BHV-1, and BVDV (Table 3). Absorbance was measured using a MultiSkan FC plate reader (Fisher Scientific, Leicestershire, UK). For each sample, one of the duplicate aliquots was heat treated at 56°C for two hours and archived at −80°C until it was shipped, on dry ice, to MRI or University of Glasgow, UK, for testing. All or a random selection of available samples were tested by indirect ELISA for *N. caninum* and *T. gondii*, and *C. abortus* (goats and sheep only) as described previously (Helmick et al., 2002, Marques et al., 2012, Wilson et al., 2009). A random selection of cross-sectional sera was also tested for exposure BTV (Table 3).

**Table 3:** Cross-sectional and abortion cohort serological tests (ELISA) for abortigenic pathogens used for various livestock species, northern Tanzania, 2013-2019.

| Target pathogen | Livestock species tested |  |  | ELISA kit name (manufacturer or source) |
| --- | --- | --- | --- | --- |
|  | Cattle | Goat | Sheep |  |
| *Brucella* spp. | ✓ | ✓ | ✓ | *Brucella* COMPELISA (Animal and Plant Health Agency, UK) |
| *Chlamydia abortus* | NA | ✓ | ✓ | In-house MRI ELISA<sup>86</sup> |
| *Coxiella burnetii* | ✓ | ✓ | ✓ | PrioCHECK Ruminant Q Fever Ab Plate § (Thermofisher®) |
| *Leptospira* Hardjo | ✓ | NA | NA | Bovine *Leptospira* Hardjo 5 Plate Solid ELISA kit (Linnodee®) |
| *Neospora caninum* | ✓ | ✓ | ✓ | In-house MRI ELISA<sup>60</sup><br>**OR**<br>*Neospora caninum* Indirect ELISA (ID Screen®) |
| *Toxoplasma gondii* | ✓ | ✓ | ✓ | In-house MRI ELISA<sup>87</sup><br>**OR**<br>Toxoplasmosis Indirect Multi-species ELISA (ID Screen®) |
| Bluetongue virus (BTV) | ✓ | ✓ | ✓ | Bluetongue competition kit (ID Screen®) |
| Bovine herpesvirus (BHV-1) | ✓ | NA | NA | IBR Individual Ab Test (IDEXX®) |
| Pestiviruses (BVDV/ BDV) | ✓ | NA | NA | BVDV p80 Ab ELISA (IDEXX®) |
| Rift Valley fever virus (RVFV) | ✓ | ✓ | ✓ | Rift Valley Fever Competition Multispecies ELISA kit (ID Screen®)<br>**OR**<br>In-house nucleocapsid-based ELISA pre-screen of goat and sheep CS sera (University of Glasgow) (Nyarobi, 2020) |

§: Formerly known as LSI/Vet Ruminant Q fever Serum/Milk
KCRI = Kilimanjaro Clinical Research Institute, Moshi, Tanzania; MRI = Moredun Research Institute, UK NA = not applicable, test not done. For *Leptospira*, BHV-1 and BVDV, serological testing of goat and sheep sera was not performed because the ELISA kits were not validated for small ruminant sera.

Clopper-Pearson 95% confidence intervals were calculated using the *binom* package ^85^ in R statistical software (version 3.6.0).

### Abortion cohort study

A prospective abortion cohort study was undertaken from October 2017 through September 2019 in 13 wards in Arusha, Kilimanjaro, and Manyara Regions of northern Tanzania (Fig. 1). Study wards were selected from those included in the cross-sectional exposure studies. Two wards were selected purposively from among those included in these cross-sectional studies because of existing relationships with the livestock-keeping community that were expected to promote participation. The remaining eleven wards were selected randomly from the wards targeted in the cross-sectional studies.

Livestock field officers (LFOs), animal health professionals working for the Tanzanian Ministry of Livestock and Fisheries, were recruited to participate in the study with one LFO responsible for a single ward. At the start of the study the LFOs attended a training course on causes of livestock abortion, as well as the practical investigation and management of abortion events. Training included the safe collection, storage, and transport of biological samples from the aborting dam, foetus, and placental material, the use of personal protective equipment for the safe handling and disposal of aborted material and methods for data collection. Following the training course, LFOs were asked to disseminate information about the study to the community living within their respective wards, requesting livestock owners to report any instances of livestock abortion to them so that an investigation could be carried out. Following receipt of a report, the LFO was asked to pass on the information to the research coordinator, a veterinarian, for investigation. For the purposes of this study, an abortion event was defined as: farmer reported visual evidence of premature foetal loss or stillbirth in cattle, goats, or sheep. The abortion was considered eligible for inclusion in this study if the research field team or LFO could attend within 72 hours of the event occurring. Two investigation options were available. Where LFOs carried out the investigation, vaginal swabs and 10 ml blood samples were collected in red-top vacutainer tubes (Becton, Dickinson and Company, New Jersey, USA) from the dam(s) that had aborted, as identified by the livestock keeper. Where the veterinarian-led research team was able to join the LFO’s within 72 hours, additional samples were collected, including swabs from the surface of the foetus exterior and placental (cotyledon) tissues, when available.

Duplicate swab samples (breakpoint shaft, viscose tip; Technical Service Consultants Ltd, Lancashire, UK) and duplicate tissue samples were stored in 1 ml DNA/RNA shield (Zymo Research, California, USA) to inactivate infectious agents and preserve nucleic acid integrity prior to molecular testing. Due to practical limitations, samples comprehensive sets of tissues including foetal fluids for bacterial culture were not collected in this study. Tissue samples were also stored in duplicate in 10% buffered formal saline to preserve tissue structure for histopathology and IHC. Samples were transported to the Zoonoses Laboratory at KCRI, for archiving at −80°C prior to testing against a panel of abortigenic agents.

### Abortion cohort serological testing

Sera from dams that aborted were tested using the range of host and pathogen specific assays described in Table 3. For each sample, one of these duplicate tubes was heat treated at 56°C for two hours prior to export to the UK for testing. The remaining tube was retained for testing at KCRI and stored at −80°C. Households were re-visited four to six weeks later to collect a convalescent blood sample from the same dam(s). Seroconversion was defined as a negative acute serum with a positive convalescent serum.

### DNA and RNA extraction

RNA was extracted using a RNeasy® Mini kit (QIAGEN, Hilden, Germany) following manufacturer’s instructions. For RNA extraction from the placental tissues, a sterile scalpel was used to add approximately 25 mg of tissue to 600 µl RLT buffer in a sterile bead beating tube containing ceramic beads (CK28, Precellys, Bertin Instruments, France). Bead beating was performed for one minute at 3,450 rpm using a MiniBeadBeater-16 (BioSpec Products, Inc., Oklahoma, USA). The liquid was aliquoted into a microcentrifuge tube with 590 µl nuclease-free water and 10 µl Proteinase K (QIAGEN, Hilden, Germany). For RNA extraction from swabs, 200 µl sample liquid was added to 600 µl RLT buffer, and 10 µl Proteinase K. All tubes were then incubated at 55°C for 10 min. Once cooled to room temperature, tubes were centrifuged at 10,000 x g for three minutes. The supernatant was transferred to a new microcentrifuge tube where half the volume of ethanol (96-99%) was added to the tube (e.g., 450 µl ethanol to 900 µl supernatant) and mixed by pipetting up and down gently. RNA was extracted from 700 µl of the lysate/ethanol mix using a final elution volume of 50 µl.

DNA was extracted using a DNeasy® blood & tissue kit (QIAGEN), following manufacturer’s instructions. For DNA extraction from placental tissues, a sterile scalpel was used to add approximately 25 mg of placenta into a microcentrifuge tube with 180 µl ATL buffer and 20 µl Proteinase K. The tubes were incubated at 56°C, vortexing occasionally, until the tissue was lysed, after which, 200 µl AL buffer was added, and the tube was vortexed briefly. For DNA extraction from swabs, 200 µl sample liquid was added to 200 µl AL buffer with 20 µl Proteinase K and vortexed. All tubes were incubated at 56°C for 10 min. When the buffer/sample mix had cooled to room temperature, 200 µl ethanol (96-99%) was added, and the tube was vortexed. DNA was extracted from the entire volume of the lysate/ethanol mix. Final elution volumes for swabs and placental tissue samples were 100 µl and 200 µl, respectively.

All batches of nucleic acid extractions included a negative extraction control where sample was replaced by nuclease-free water and the run alongside the nucleic acid extraction process.

### Molecular testing

The molecular targets for all pathogens are listed in Table 4. Primer and probe sequences and cycling conditions for each PCR, qPCR, and RT-qPCR assay, are detailed in supplementary materials (S1; Table C). DNA or RNA extracts from positive controls listed in Table 4 were used for individual PCR, qPCR, and RT-qPCR assays. All available vaginal swab, foetal swab, and placental tissue samples for all three livestock species were for all pathogens included in this study. Samples were tested in duplicate except for pathogens tested using commercial kits (BTV and pestiviruses) where single samples were tested. Nuclease-free water and extraction controls were included in each PCR, qPCR, and RT-PCR run as negative controls.

**Table 4:**
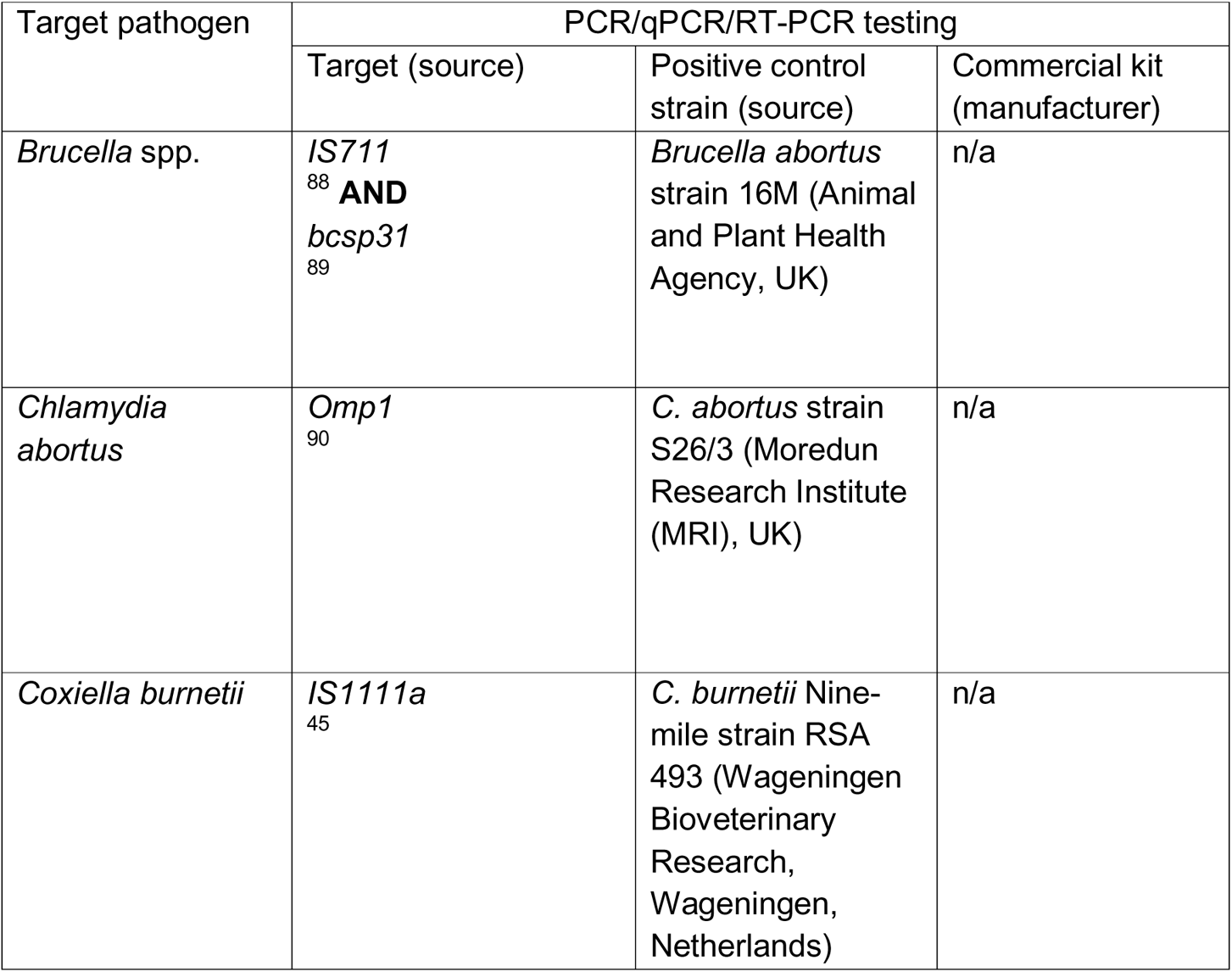
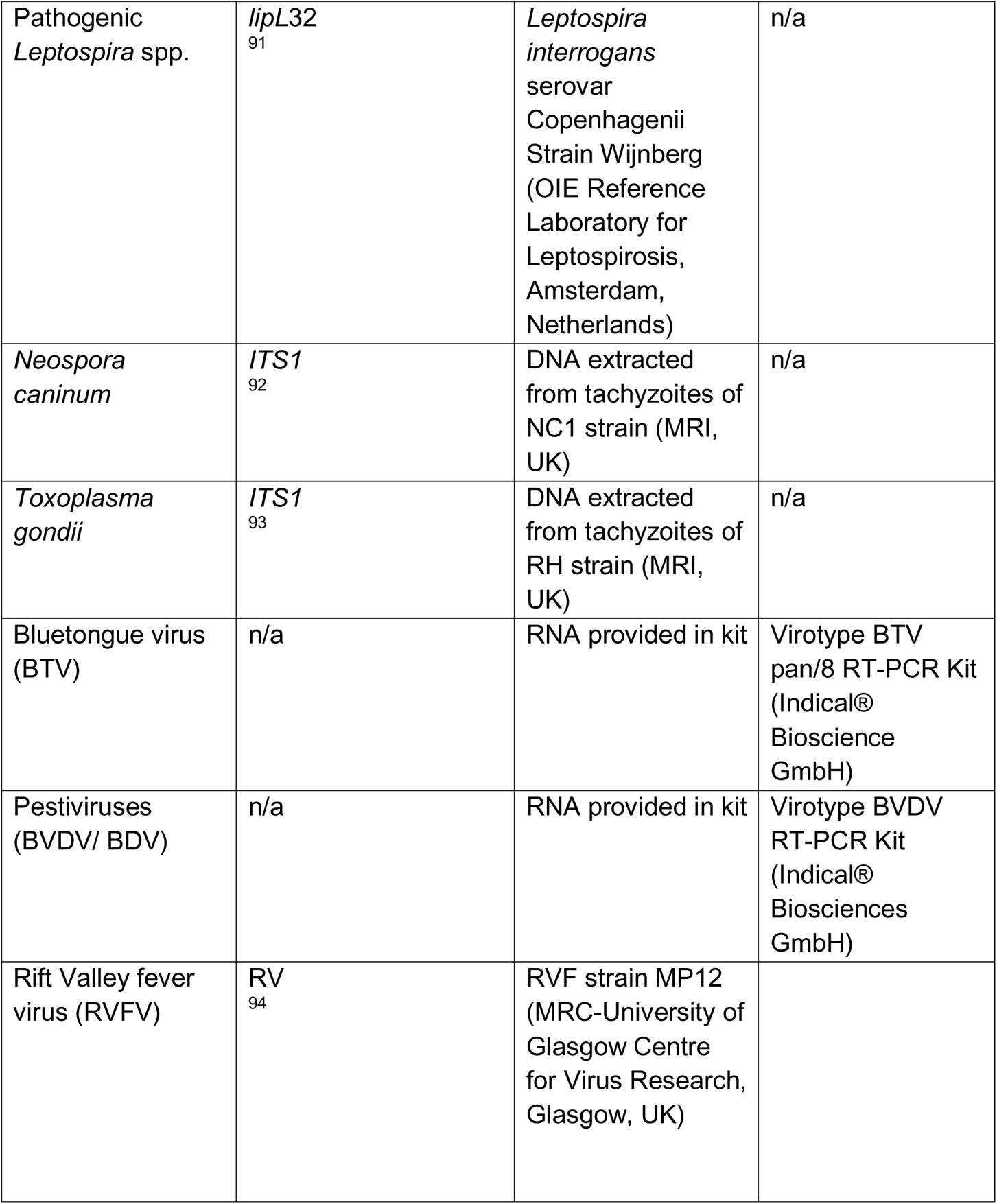
Abortion cohort study molecular assays (PCR, qPCR and RT-qPCR) for cattle, goat and sheep samples for each target pathogen, northern Tanzania, 2017-2019.

Samples were considered positive only if negative extraction controls and negative controls showed no evidence of amplification and the internal and external positive controls also amplified. Where single wells amplified for samples tested in duplicate, the assay was repeated. Unless specified, testing was performed at KCRI.

Previously published qPCR assays designed and validated for the diagnosis of infection with the respective pathogens were used to detect the presence of *Brucella* spp. (two gene targets ^88, 89^), *C. burnetii* ^45^, *C. abortus* ^90^, and pathogenic *Leptospira* spp. ^91^. These qPCRs were performed in single well reactions using a QuantiNova® Probe kit (QIAGEN) with 5 μl of DNA template (or 2.5 μl per target for *Brucella* spp.).

Nested conventional PCR assays for the presence of *Neospora* spp. and *Toxoplasma gondii* were performed targeting the internal transcribed spacer (ITS1) region between the 18S and 5.8S rRNA genes ^95^. The primary (1°) nested PCR was performed using 2 μl of DNA template. The secondary (2°) nested PCR was performed using with 2 μl of a 1:5 diluted primary PCR product. Both 1° and 2° assays were performed with 2x PCR master mix (Promega Corporation, Wisconsin, USA). The final PCR product from the 2° nested PCR was checked by electrophoresis in a 1.5% agarose gel. Samples were considered positive if the size of bands visible under UV light were congruous with those expected of the *Neospora* (249 bp) and *Toxoplasma* (227 bp) targets.

The presence of BTV and pestivirus RNA in samples was determined by RT-qPCR using commercial kit-based assays validated for the detection of the respective pathogens (Table 4). Single samples were tested, as per kit instructions, using 5 μl of RNA template. Finally, the presence of RVFV RNA was assessed using the QuantiNova® RT-PCR probe kit (QIAGEN). Samples were tested in duplicate using 5 μl of RNA template per well.

### Pestivirus speciation

As commercial kit-based RT-qPCR assays for pestiviruses do not distinguish between the different pestivirus species, genotyping was performed at MRI to determine the infecting pestivirus species. The RNA extracts that were positive in pan-pestivirus RT-PCR screening assays were subjected to RT-PCR and then nested PCR using four primer pairs developed for genotyping of livestock pestiviruses: 324-326 (5’UTR); BD1-BD3/BD4 (N^pro^); BD1-BD2 (N^pro^-C); and 17f-1400r (UTR-Erns) to determine the infecting virus type ^96–98^. RT-PCR products of the expected sizes were purified from agarose gels and submitted for sequencing using the relevant PCR primers (Eurofins-MWG). Sequence traces for each sample were assembled using SeqMan Pro (DNASTAR Lasergene) and consensus sequences that excluded the terminal primer sites were generated. The 504 nt N^pro^ coding sequences were aligned with a set of N^Pro^ reference sequences that included representatives of pestivirus species A-H plus K, as described previously ^36^.

Alignment was performed using MAFFT (https://mafft.cbrc.jp/alignment/). Maximum likelihood (ML) phylogenetic analysis was performed using MEGA version X. Model selection was used to identify the best evolutionary model for the dataset. The ML analysis was bootstrapped 500 times to estimate the reliability of the resulting phylogeny.

### Additional pathogen genotyping

*T. gondii* genotyping was also performed to determine the infecting sub-type. The methods of this genotyping work are detailed in the supplementary materials S1; D.

### Pathology and immunohistochemistry (IHC)

Histopathology was performed on placental tissue samples where available. For samples that tested positive for *C. burnetii* by qPCR, *C. burnetii* IHC was also performed. The methods for the histopathology and IHC are given in supplementary materials S1; B.

### Diagnosis of abortion aetiology

For the purposes of our study, we determined a set of case definitions that we used to attribute livestock abortion events to a specific aetiology. These case definitions are shown in Table 5. Case definitions were based on OIE guidelines for the diagnosis of each pathogen. Due to the challenges in systematically collecting tissue samples for histopathology from abortion cases (e.g. lack of available tissue in many cases), we designed these diagnostic criteria (Table 5) to be independent of histopathological examination but used evidence from histopathological examination to support our diagnosis where samples existed to support this diagnostic approach. For *N. caninum*, for which no OIE diagnostic guidelines exist, the case definition for was based on criteria used by MRI, UK ^99^, a specialist diagnostic veterinary laboratory. For *C. burnetii,* OIE guidelines indicate that definitive diagnosis of infectious abortions resulting from *C. burnetii* infection can only be robustly achieved at the flock or herd level, but also that bacterial load is predictive of causality in individual animals ^79, 80^. Therefore, to determine the *C. burnetii* case definitions, we used a two-step approach. First, we used the minimum bacterial load associated with abortions as stated by the OIE (> 10^4^ bacteria per swab or per gram of tissue, equivalent to Ct values of 37.5 and 42.7 respectively on our PCR platform).

**Table 5:** Diagnostic criteria used by this study to attribute the aetiology of abortions in cattle, sheep, and goats reported to the abortion surveillance platform. Diagnostic case definitions apply to all three ruminant species unless otherwise specified.

| Pathogen | Were OIE diagnostic guidelines available? | Summary of OIE diagnostic criteria<br>(presuming recent history of abortion in all cases) | Diagnostic case definition applied in this study | Confidence in diagnostic case definition |
| --- | --- | --- | --- | --- |
| *Brucella* spp. | Yes - for aetiological diagnosis of abortion<sup>100</sup> | <u>**Definitive diagnosis (dx):**</u> Isolation of *Brucella* from abortion material, udder secretions or from tissues removed at post-mortem<br><u>**Presumptive diagnosis (dx):**</u> Evidence of *Brucella* by the demonstration of *Brucella*-like organisms in abortion material or vaginal discharge using modified acid-fast staining, if supported by serological tests. Polymerase chain reaction (PCR) methods are additional means for detection of the presence of *Brucella* DNA in a sample. | Vaginal swab, foetal swab and/or placental tissue samples positive by qPCR against both *IS711* (Ct <35) and *bcsp31* (Ct <40) targets<br>**AND**<br>Evidence of seropositivity in acute and/or convalescent serum samples from dam | Presumptive |
| *Chlamydia abortus* | Yes - for aetiological diagnosis of abortion in small ruminants<sup>101</sup> | <u>**Definitive dx:**</u> Evidence of purulent to necrotising placentitis with vasculitis by histopathology, and demonstration of large number of the organism in placenta or vaginal excretions by PCR or antigen tests or stained smears.<br><u>**Presumptive dx:**</u> In the absence of samples for direct pathogen detection, a rise in | Vaginal swab, foetal swab and/or placental tissue samples positive by qPCR (Ct < 40)<br>**OR**<br>In the absence of a vaginal swab or placental sample, evidence of seroconversion | Definitive<br>Suggestive |
|  |  | *antibody titre (detected by ELISA) but does not occur in every case.* | between acute and convalescent serum samples from dam (goats and sheep only) |  |
| *Coxiella burnetii* | Yes - for aetiological diagnosis of abortion at the herd or flock level only<sup>80</sup> | <u>Presumptive dx (individual animal):</u><br>*Positive identification of high levels of C. burnetii >10<sup>4</sup> per vaginal swab or per gram of placental tissue<sup>79, 102</sup> by PCR in placenta or vaginal discharges.* | Placental tissue, vaginal swab or foetal swab positive by qPCR by IS1111a qPCR (Ct ≤ 27)* | Presumptive |
| Pathogenic *Leptospira* spp. | Yes - for diagnosis of dam infection only<sup>103</sup> | <u>Definitive dx:</u> Isolation of *Leptospira* (recommended) or demonstration by PCR (suitable method) from blood or milk<br><u>Presumptive dx:</u> demonstration of agent (isolation or PCR) in the genital tract or vaginal fluids in association with compatible clinical signs and evidence of serological exposure | Vaginal swab, foetal swab and/or placental tissue samples positive by qPCR (Ct < 40)<br>**OR**<br>Evidence of seroconversion between acute and convalescent serum samples from dam (cattle only)<sup>59</sup> | Presumptive |
| *Neospora caninum* | No | N/A | Vaginal swab, foetal swab and/or placental tissue samples positive by PCR<sup>104</sup> | Presumptive |
| *Toxoplasma gondii* | Yes - for aetiological diagnosis of abortion<sup>105</sup> | <u>Definitive dx:</u> Demonstration of *Toxoplasma* in the placenta of an aborted foetus by either histopathology or PCR | Vaginal swab, foetal swab and/or placental tissue samples positive by PCR<sup>53</sup> | Definitive |
| Bovine herpes virus-1 (BHV-1) | Yes - for aetiological diagnosis of abortion<sup>106</sup> | <u>Definitive dx:</u><br>*Demonstration of BHV-1 by PCR or virus isolation from abortion material.*<br><u>Presumptive dx:</u><br>*Demonstration of BHV-1 by PCR or virus isolation from swabs from the genital tract or evidence of seroconversion between acute and convalescent samples confirming acute infection demonstrated by ELISA or virus neutralisation.* | Evidence of seroconversion between acute and convalescent serum samples from dam (cattle only) | Suggestive |
| Bluetongue virus (BTV) | Yes - for aetiological diagnosis of abortion<sup>107, 108</sup> | <u>Definitive dx:</u> *Detection of pathogen by RT-PCR or classical virus isolation in blood or tissue samples including from aborted or stillborn animals.* | Vaginal swab, foetal swab and/or placental tissue samples positive by RT-qPCR (Ct < 40) | Definitive |
| Pestiviruses (BVDV or BDV) | Yes - for aetiological diagnosis of abortion<sup>109</sup> | <u>Definitive dx:</u> *Detection of pathogen by RT-PCR, virus isolation or immunohistochemistry in foetal tissues or fluids.* | Vaginal swab, foetal swab and/or placental tissue samples positive by RT-qPCR (Ct < 40) | Definitive |
| Rift Valley fever virus (RVFV) | Yes - for aetiological diagnosis of abortion<sup>110</sup> | <u>Definitive dx:</u> *Requires a combination of at least two positive test results from two diagnostic test methods including detection of pathogen by virus isolation or RT-PCR performed on blood (from dam), tissue from aborted foetuses or other abortion products; positive detection of antibodies (serum); or* | Vaginal swab, foetal swab and/or placental tissue samples positive by RT-qPCR (Ct < 40) **AND** Detection of RVFV IgG in acute and/or convalescent serum from dam by RVFV | Definitive |
|  |  | *positive for IgM or IgG ELISA with demonstration of rising titres between paired serum samples collected 2-4 weeks apart.* | IgG ELISA <sup>110</sup> |  |

\*See supplementary materials S1; E for further details

Secondly, using a histogram of the Ct results at the study population-level, we identified a bimodal distribution in the results, suggesting two distinct biological processes within our sampled population (see supplementary material S1; E). We used this frequency distribution, supported by histopathology and IHC results to select a conservative threshold value using a cut-off Ct value of ≤ 27 for attributing abortion events to *C. burnetii* in this study.

Confidence in this study’s diagnostic case definition was determined to be either ‘definitive’, ‘presumptive’ or ‘suggestive’, depending on how closely our diagnostic approach aligned with the OIE guidelines for diagnosis of abortions caused by each pathogen. Where our study diagnostic approach met OIE definitions for a definitive diagnosis, we classified our diagnosis as ‘definitive’ and had the highest confidence in our aetiologic diagnosis. Where our study diagnostic approach met OIE definitions for a presumptive diagnosis or for *N. caninum* where no OIE guidelines exist, we classified our diagnosis as ‘presumptive’. Where our study showed evidence of recent dam infection by serology, but did not detect the presence of the pathogen directly in any samples (e.g., for BHV-1 in cattle, or *C. abortus* in small ruminants), we classified the diagnosis as ‘suggestive’ indicating a lower level of confidence in these aetiologic diagnoses.

## Supplementary materials

**A: Table A**: Cross-sectional and abortion cohort acute serology

**B:** Histopathology and immunohistochemistry for abortion cohort study including **Table S1B** (Summary of results from histopathologic examination of placental samples collected from aborting ruminant livestock as part of the abortion cohort survey, northern Tanzania) and interpretation and conclusions.

**C: Table C**: Target pathogen serologic (ELISA) and molecular (PCR/RT-PCR) assay information, including primer/probe sequences and cycling conditions for samples from various livestock species.

**D:** Genetic characterisation of *T. gondii* isolated from an abortion case

**E:** Determining qPCR cut-off values for diagnosing *Coxiella burnetii* as the cause of abortion in ruminant livestock in Tanzania.

## Acknowledgements

The abortion cohort study was supported by the Supporting Evidence Based Interventions project, University of Edinburgh (grant number R83537 CH). The cross-sectional studies were supported by US National Institutes of Health-National (NIH) Science Foundation Ecology and Evolution of Infectious Disease program (R01 TW009237) and the UK Biotechnology and Biological Sciences Research Council (BBSRC) (grant numbers: BB/J010367/1, BB/L018926/1). MD was funded by the Scottish Government. CH, CH, EI, FK, ML, GR and DL are supported by funding from the Scottish Government Rural and Environment Science and Analytical Services Division (RESAS). BW is funded by the BBSRC. We would like to thank Rosanne de Jong, Divine Ekwem, Erin Hodgkinson, Karen Main and Jen Jen Yu for assistance with serological testing at the University of Glasgow and Clare Underwood (MRI) for expert immunohistological preparations. We appreciate the huge efforts of our field team, Rigobert Tarimo, Fadhili Mshana, and Hassan Hussein as well as the assistance from District Veterinary Officers and LFOs. Thanks to Paul Johnson (University of Glasgow) for support with data management and curation, and Dassa Nkini (NM-AIST) and Elizabeth Kussaga (KCRI) for administrative support.

## Author contributions

Developed the study concept and design of the work: KT, JRC, WdG, FL, JB, JAC, MD, JH, CH, EAI, FK, DL, BM, ON, NW, SC, KA; Collected samples and metadata: TK, WdG, FL, GS; Sample processing KT, NA, RC, GC, MD, CH, EI, FK, ML, CM, VM, JN, GR, NW, BW, SC, KA; Data analysis and interpretation: KT, JRC, WdG, FL, NA, RC, GC, JAC, MD, JH, CH, EI, FK, ML, DL, CM, VM, JN, GR, NW, BW, SC, KA; Initial drafting the article: KT, WdG, FL, JH, SC, KA; All authors provided critical revision of the article and provided approval for submission.

## Competing interests

The authors declare no competing interests.

## Notes

### Competing Interest Statement

The authors have declared no competing interest.

### Summary of Updates

Reformatted with pestivirus sequence accession numbers added.

