## Supplementary Materials A-E for "Prospective cohort study reveals unexpected aetiologies of livestock abortion in northern Tanzania" (from another version of this preprint: version 1, 2021-08-26)

Supplementary Materials (S1A-E) for: **Prospective cohort study reveals unexpected aetiologies of livestock abortion in northern Tanzania**

**S1, A; Table A: Individual-level prevalence (% seropositive) of exposure to a range of abortigenic agents in livestock in northern Tanzania sampled through: (i) cross-sectional surveillance and (ii) abortion cohort study (acute serum samples only).** Clopper-Pearson 95% confidence intervals are also shown [ ].

|  | **Cattle** | | | **Goats** | | | **Sheep** | | |
| --- | --- | --- | --- | --- | --- | --- | --- | --- | --- |
| **Abortigenic agent** | **Cross-sectional** | | **Abortion cohort (n = 71)** | **Cross-sectional** | | **Abortion cohort (n = 98)** | **Cross-sectional** | | **Abortion cohort (n = 43)** |
| **Bacteria** | **n tested** | **%** | **%** | **n tested** | **%** | **%** | **n tested** | **%** | **%** |
| *Brucella* spp. | 3627 | 4.1 [3.5 – 4.8] | 5.6 [1.6 – 13.8] | 3342 | 3.7 [3.1 – 4.4] | 0 [0 – 3.7] | 2628 | 2.5 [2.0 – 3.2] | 0 [0 – 8.2] |
| *Chlamydia abortus* | n/a | n/a | n/a | 1429 | 1.3 [0.8 – 2.0] | 1.1 [0.0 – 6.0]^a^ | 1017 | 0.8 [0.57 – 3.1] | 2.4 [0.1 – 12.6]^b^ |
| *Coxiella burnetii* | 3602 | 5.2 [4.5 – 6.0] | 2.8 [0.3 – 9.8] | 3325 | 27.9 [26.3– 29.4] | 37.8 [28.2 – 48.1] | 2620 | 17.4 [15.9 – 18.9] | 20.9 [10.0 – 36.0] |
| *Leptospira* Hardjo serovars | 3527 | 22.3 [21.0 – 23.8] | 8.5 [3.2 – 17.5] | n/a | n/a | n/a | n/a | n/a | n/a |
| **Protozoa** |  |  |  |  |  |  |  |  |  |
| *Neospora caninum* | 3108 | 12.6 [11.4 – 13.8] | 31.0 [20.5 – 43.1] | 479 | 1.7 [0.7 – 3.3] | 1.0 [0.0 – 5.6]^c^ | 412 | 1.5 [0.5 – 3.1] | 0 [0 – 8.2] |
| *Toxoplasma gondii* | n/a | n/a | 18.3 [10.1 – 29.2] | 1420 | 14.9 [13.1 – 16.8] | 23.7 [15.7 – 33.4]^c^ | 1009 | 10.6 [8.8 – 23.7] | 7.0 [1.5 – 19.1] |
| **Viruses** |  |  |  |  |  |  |  |  |  |
| Bovine Herpes Virus 1 (BHV-1) | 3013 | 66.2 [64.5 – 67.9] | 28.2 [18.1 – 40.1] | n/a | n/a | n/a | n/a | n/a | n/a |
| Bluetongue virus (BTV) | 299 | 90.3 [86.4 – 93.4] | n/a | 186 | 86.0 [80.2 – 90.7] | n/a | 367 | 68.7 [63.7 – 73.4] | n/a |
| Bovine viral diarrhoea virus (BVDV) | 3015 | 36.0 [34.3 – 37.8] | 49.3 [37.2 – 61.4] | n/a | n/a | n/a | n/a | n/a | n/a |
| Rift Valley fever virus (RVFV) | 3582 | 4.4 [3.7 – 5.1] | 21.1 [12.3 – 32.4] | 3304 | 1.4 [1.0 – 1.8] | 2.0 [0.2 – 7.2] | 2583 | 2.6 [2.0 – 3.3] | 0 [0 – 8.2] |

n/a = not applicable - tests not run; (a) n = 90 for *C. abortus* testing in goats; (b) n = 42 for *C. abortus* testing in sheep; (c) n = 97 for *N. caninum* and *T. gondii* testing in goats

**S1, B: Histopathology and immunohistochemistry for abortion cohort study**

**Objective:** To perform histopathologic examination and targeted immunohistochemistry on available placental tissues to support diagnosis of aetiology of abortions from ruminant livestock.

**Methods:**

Abortion events were identified as outlined in the main text of this article. Following an abortion event, where placental tissue samples were available, sections of cotyledonary tissue was stored in 10% buffered formal saline to preserve tissue structure for histopathology and immunohistochemistry (IHC). Formalin-fixed tissue samples were shipped to University of Glasgow, UK under import licenses TARP(S) 2019/07 and Material Transfer Agreements between Kilimanjaro Christian Medical Centre (KCMC) and University of Glasgow, and between KCMC and Moredun Research Institute (MRI), Edinburgh, Scotland.

Histopathology was performed at the University of Glasgow Veterinary Diagnostic Services (UG VDS) with examination conducted by two veterinary pathologists (CM and GC). Available tissue samples were processed routinely and sections (4 μm) were stained with haematoxylin and eosin (HE). Special stains were applied at the discretion of the examining pathologist following standard methods including Gram stains, Giemsa, and Periodic Acid Schiff (PAS) to diagnose the presence of bacteria, fungi, or other micro-organisms. Finally, Von Kossa stain was performed to differentiate mineral deposits, a normal histologic finding in ruminant placenta (Botta *et al.*, 2019), from other causes of granular deposits (e.g., bacteria) within placental tissue.

Subsequently, placental tissue samples that tested positive for *Coxiella burnetii* by *IS1111a* qPCR on any sample were transported to MRI for IHC. Sections of placental tissue were deparaffinized in xylene prior to rehydration to 95% ethanol. After quenching of endogenous tissue peroxidase activity with 3% hydrogen peroxide (Sigma) in methanol (v/v), the sections were washed in water followed by Tris Buffered Saline (TBS). Non-specific antibody binding was blocked with 25% normal goat serum (NGS, Vector Laboratories, Peterborough, UK) in TBS for 30 min prior to overnight incubation at 4 °C with primary mouse monoclonal anti-*C. burnetii* antibodies (mAb) (clone 3.13; Squarix GmbH, Marl, Germany), diluted in 1/500 in TBS. Visualisation of bound antibodies was by the EnVision Kit (goat anti-mouse horseradish peroxidase conjugate, DakoCytomation, Ely, UK), as per the manufacturer’s instructions, followed by addition of chromogen (AEC, Vector Laboratories, Peterborough, United Kingdom) for 10 min. Slides were then washed with tap water, counterstained with haematoxylin for 1 min, washed in tap water again and mounted in ImmunoHistoMount™ (Sigma-Aldrich Company Ltd., Poole, United Kingdom). Negative control sections were comprised of semi-serial sections of all tissues examined by IHC, with the primary antibodies substituted by isotype-matched mouse IgG antibodies. Sections of known *C. burnetii*-positive bovine placenta from a clinical case of Q-fever abortion were also included as positive controls.

**Results:**

Fixed placental tissue was available for histopathology from 41 (19.1%) of 215 abortion events recorded by the study. By species, this included 32 (45.1%) of 71 cattle abortion events, eight (8.0%) of 100 goat abortion events and one (2.3%) of 44 sheep abortion events. Of abortion events with placental tissue available for examination, 18 (43.9%) met the study case definitions as defined in Table 3 (main text). *Neospora caninum* was the most commonly diagnosed pathogen in eight (25.0%) of 32 cattle abortion events and one (12.5%) of eight goat events with placental tissue available. In addition, Rift Valley fever virus (RVFV) was diagnosed in six (18.8%) of 32 cattle events, bovine herpesvirus (BHV-1) in two (6.3%) cattle, bovine viral diarrhoea virus (BVDV) in one (3.1%) cow and *C. burnetii* in one (12.5%) goat. Of note, one cow met the case definition for both *N. caninum* and BHV-1.

A summary of histopathology findings is shown in Table S1C. Autolysis, ranging from mild to severe, or poor tissue preservation was observed as a feature in placental samples from 38 (92.7%) of 41 abortion events, which limited histopathological interpretation in some cases. Evidence of placental inflammation (placentitis) was the most common pathological lesion identified and was detected in 14 (34.1%) abortion events, from 13 (40.6%) cattle and one (12.5%) goat. Vasculitis also accompanied the placentitis in eight (25.0%) cattle abortion events. Accumulations of bacteria within placental tissue were a confirmed by Gram stain in 11 (84.6%) of 13 abortion events where Gram staining was performed. In five (45.4%) of these, the bacteria present were of mixed morphology and were interpreted as most likely to be a result of post-mortem contamination. The significance of other bacteria observed is unclear without further bacteriological or metagenomic investigation.

Positive antibody labelling for *C. burnetii* was detected in the placenta of one goat that met our study case definitions for *C. burnetii-*associated abortion. A further seven animals (five cattle, one goat, one sheep) that were positive for *C. burnetii* by *IS1111a* qPCR, but did not meet our study case definitions were also subjected to IHC but no significant antibody labelling, or pathology indicative of *C. burnetii* infection, was detected in any of these samples.

**Table S1, B: Summary of results from histopathologic examination of placental samples collected from aborting ruminant livestock as part of the abortion cohort survey, northern Tanzania.**

| **Animal ID** | **Species** | **Tissue preservation** | **Special stains & results** | **Coxiella IHC** | **Morphological Diagnosis** | **Pathogens detected by PCR** | | | **Evidence of seroconversion (paired serology)** | **Diagnosis based on study case definitions** |
| --- | --- | --- | --- | --- | --- | --- | --- | --- | --- | --- |
|  |  |  |  |  |  | **Placenta** | **Vaginal swab** | **Foetal swab** |  |  |
| SEBI-010 | Cattle | Marked autolysis | Gram stain: No significant findings | NA | None | No | *Neospora caninum* | *Neospora caninum* | *Neospora caninum* | *Neospora caninum* |
| SEBI-011 | Cattle | Poor tissue preservation | Gram stain: Moderate to high numbers of gram positive and Gram-negative bacteria including large aggregates of gram negative and separately Gram-positive coccobacilli.    PAS stain: no significant findings | NA | Placentitis, suppurative, lymphohistiocytic | No | *Neospora caninum* | *Neospora caninum* | *Neospora caninum* | *Neospora caninum* |
| SEBI-013 | Cattle | Moderate autolysis, poor tissue preservation | Gram stain: Multifocally low numbers of Gram-positive bacilli are seen scattered and in rare clusters within the tissue    PAS stain: Positive indicating presence of glycos-aminoglycans    Von Kossa: Negative | NA | Placentitis, suppurative | No | No | NA | BHV-1 | BHV-1 |
| SEBI-016 | Cattle | Some autolysis | NA | NA | None | *Neospora caninum* | No | No | *Neospora caninum* | *Neospora caninum* |
| SEBI-028 | Cattle | Marked to severe autolysis | Gram stain: Moderate to high numbers of gram negative coccibacilli within tissue, and layer of Gram-positive cocci bacilli and few rods adherent to the surface of the tissue. | NA | Placentitis, suppurative and histiocytic, with necrotising vasculitis, and multifocal fibrin thrombi | No | No | No | No | None |
| SEBI-029 | Cattle | Moderate to marked autolysis | NA | NA | None | *Neospora caninum* | No | No | *Neospora caninum* | *Neospora caninum* |
| SEBI-044 | Cattle | Marked autolysis | NA | NA | None | No | No | No | No | None |
| SEBI-046 | Cattle | Moderate to marked autolysis | Gram stain: No significant findings    PAS stain: No significant findings | No labelling | None | *Coxiella burnetii*(Av Ct 36.0) | No | No | No | None |
| SEBI-054 | Goat | No significant preservation problems reported | Gram stain: Few trophoblasts in the area of necrosis have cytoplasm which stains Gram negative, but no distinct bacteria can be seen    PAS stain: Low to moderate number of trophoblasts contain moderate amounts of PAS positive finely granular material (1um in size) | Variably small to medium amounts of positive granular labelling present within the cytoplasm of small to moderate numbers of trophoblasts | Placentitis, necrotising, suppurative | *Coxiella burnetii*  (Av Ct 12.5) | *Coxiella burnetii* (Av Ct 9.3) | NA | NA | *Coxiella burnetii* |
| SEBI-055 | Cattle | No significant preservation problems reported | Gram stain: Very low scattered numbers of Gram-positive rods are seen within autolytic cotyledons | A very small amount of granular labelling within the cytoplasm of a low number of degenerate cells plus a large amount of very pale and diffuse labelling | None | *Coxiella burnetii*  (Av Ct 32.2)    *Neospora caninum* | *Coxiella burnetii*  (Av Ct 30.2) | NA | *Neospora caninum* | *Neospora caninum* |
| SEBI-067 | Goat | Mild to moderate autolysis | NA | NA | None | No | No | No | No | None |
| SEBI-076 | Cattle | Marked autolysis | NA | No labelling | None | *Coxiella burnetii*  (Av Ct 36.7)    RVFV | *Coxiella burnetii*  (Av Ct 36.5)    RVFV | RVFV | NA | RVFV |
| SEBI-079 | Cattle | Moderate to marked autolysis | NA | NA | None | RVFV | No | No | No | RVFV |
| SEBI-084 | Cattle | Moderate to marked autolysis | NA | NA | None | RVFV | No | RVFV | No | RVFV |
| SEBI-097 | Goat | Mild autolysis | NA | NA | None | No | No | NA | No | None |
| SEBI-099 | Cattle | Moderate to marked autolysis | NA | NA | None | RVFV | No | NA | No | RVFV |
| SEBI-102 | Goat | Mild to marked autolysis | NA | NA | None | No | No | NA | NA | None |
| SEBI-103 | Cattle | Moderate to marked autolysis | Gram stain: Moderate to high numbers of Gram-negative coccobacilli/short rods (also stain positive with Giemsa) seen in tissue and in adjacent debris on surface    PAS stain: No significant findings | NA | Placentitis, lymphohistiocytic, with necrotising vasculitis and high numbers of Gram negative, Giemsa positive bacteria | No | No | NA | No | None |
| SEBI-110 | Cattle | Moderate to marked autolysis | NA | NA | None | RVFV | RVFV | RVFV | No | RVFV |
| SEBI-114 | Cattle | Marked autolysis | NA | NA | None | No | No | NA | No | None |
| SEBI-115 | Cattle | Marked autolysis | NA | No labelling | None | *Coxiella burnetii* (Av Ct 36.7)    RVFV | RVFV | NA | No | RVFV |
| SEBI-116 | Cattle | Variable preservation with moderate to marked autolysis in some sections and reasonable preservation in other areas | Gram stain: Low numbers of Gram-negative bacteria seen multifocally (Giemsa negative)  PAS stain: No significant findings | NA | Placentitis, suppurative, necrotising with necrotising vasculitis and fibrin thrombi. | No | No | No | No | None |
| SEBI-117 | Cattle | Variable moderate to severe autolysis | NA | NA | Placentitis, lymphohistiocytic with necrotising vasculitis | No | No | NA | No | None |
| SEBI-122 | Cattle | Marked autolysis | NA | NA | None | No | No | NA | No | None |
| SEBI-127 | Cattle | Marked to severe autolysis | NA | NA | Placentitis, suppurative | No | No | NA | No | None |
| SEBI-130 | Cattle | No significant preservation problems reported | NA | NA | Placentitis, lymphohistiocytic with vasculitis and fibrin thrombi | No | No | No | NA | None |
| SEBI-133 | Sheep | Moderate autolysis | NA | A small number of trophoblasts with small to medium amounts of positive labelling within the cytoplasm | None | No | *Coxiella burnetii*  (Av Ct 37.6) | NA | No | None |
| SEBI-135 | Cattle | Moderate to marked autolysis | Gram stain: There are low numbers of gram positive bacteria and few gram negative cocci distributed in surface debris throughout the section. Post mortem contamination suspected. | NA | Placentitis, suppurative | No | No | No | NA | None |
| SEBI-136 | Cattle | Variable preservation | NA | NA | Placentitis, lymphohistiocytic with necrotising vasculitis | No | No | NA | NA |  |
| SEBI-168 | Goat | Moderate autolysis | NA | A single cell with diffuse cytoplasmic labelling | None | *Coxiella burnetii*  (Av Ct 36.3) | *Neospora caninum* | *Neospora caninum* | *Neospora caninum* | *Neospora caninum* |
| SEBI-169 | Goat | Moderate to marked autolysis | NA | NA | None | No | No | No | No | None |
| SEBI-172 | Cattle | Moderate to marked autolysis | Gram stain: Moderate to high numbers of Gram-positive coccobacilli are present admixed with debris on the surface of cotyledons and near the surface of villi | NA | Placentitis, necrotising, suppurative | No | No | NA | No | None |
| SEBI-175 | Cattle | Moderate to marked autolysis | NA | No labelling | None | *Coxiella burnetii*  (Av Ct 37.5) | No | No | No | None |
| SEBI-184 | Cattle | Moderate to marked autolysis | Gram stain: Moderate numbers of Gram-positive and Gram-negative rods seen on the surface debris and fewer within stroma. Post mortem contamination suspected. | NA | Placentitis, necrotising, lymphohistiocytic and suppurative with necrotising vasculitis and fibrin thrombi. | *Neospora caninum* | *Neospora caninum* | NA | BHV-1  *Neospora caninum* | BHV-1 and *Neospora caninum* |
| SEBI-185 | Cattle | Mild to moderate autolysis | NA | NA | None | *Neospora caninum* | No | No | *Neospora caninum* | *Neospora caninum* |
| SEBI-186 | Goat | Severe autolysis | NA | NA | None | No | No | No | No | None |
| SEBI-187 | Cattle | Moderate to marked autolysis | Gram stain: Gram-positive and negative bacteria of mixed morphology are present throughout the tissue including within blood vessels. Post mortem contamination suspected. | NA | Placentitis, necrotising, suppurative with vasculitis | No | No | NA | No | None |
| SEBI-189 | Cattle | Moderate to marked autolysis | NA | NA | None | *Neospora caninum* |  | NA | *Neospora caninum* | *Neospora caninum* |
| SEBI-190 | Cattle | Mild autolysis | NA | NA | None | No | No | Pestivirus (BVDV) | BVDV/BDV | BVDV |
| SEBI-191 | Cattle | Marked to severe autolysis | NA | NA | None | No | No | NA | NA | None |
| SEBI-202 | Goat | Moderate autolysis | NA | NA | None | No | No | No | NA | None |

*As defined by study case definitions

**Interpretation and conclusions:**

***Neospora caninum*-associated abortions:** *N. caninum* was the most commonly diagnosed aetiology of abortion in cattle based on study case definitions but placental histopathology results were mixed. In two cases, placental pathology including placentitis and vasculitis was observed and for the remaining six cattle cases and one goat case, no evidence of placental pathology was observed. In *N. caninum-*associated abortion, placental lesions are reported to be limited in nature, with focal areas of necrosis and non-suppurative inflammation confined to the cotyledons (Dubey *et al.*, 2006). Histopathology of foetal tissues is more typically used to confirm a diagnosis of *N. caninum* but unfortunately, foetal tissues were rarely available for examination in our study. However, the presence of placental lesions is not a pre-requisite for the diagnosis of *N. caninum-*associated abortion (Dubey & Schares, 2006), and the presence of *N. caninum* nucleic acids accompanied by evidence of dam seropositivity or seroconversion (as noted in all the positive abortion events included in this study component) remains supportive of a diagnosis of *N. caninum* in most cases.

Notably, evidence of co-infections with other pathogens was observed in three *N. caninum-*positive cases. One cattle case (SEBI-184), which met study case definitions for both *N. caninum* and BoHV-1, had evidence of placentitis accompanied by necrotising vasculitis. *N. caninum* and *C. burnetii* were detected concurrently in one cattle (SEBI-055) and one goat (SEBI-168) case respectively, although neither cases met the study case definition for *C. burnetii* due to insufficient bacterial load in placental or swab samples. It is therefore feasible that the presence of other bacterial and viral pathogens may have contributed to the severity of pathology observed in these abortions. Dubey et al (2006) note that multiple factors may interact to influence the severity of foetal or placental damage in *N. caninum-*associated abortions and therefore, it remains a plausible possibility that multiple infectious agents may have contributed to the foetal and placental damage that led to negative pregnancy outcome in these cases.

**RVFV-associated abortions***:* Placental pathology was not detected in any of the six cattle abortion events that met the study case definitions for RVFV. Recent studies of RVFV-associated abortions in sheep have indicated that abortion results from severe acute pathology of the placenta including haemorrhage within the placentomes of infected dams and necrosis of the foetal trophoblasts and maternal epithelial in the cotyledonary and intercotyledonary chorioallantois (Oymans *et al.*, 2020, Odendaal *et al.*, 2020). In a set of experimental RVFV infections in sheep, all abortions occurred within 7 days post infection indicating the acute nature of these placental changes, which were often observed in the absence of any notable foetal pathology (Oymans et al., 2020). Although cattle are reported to be slightly more resistant to RVFV-associated disease than sheep, the pathology associated with infection are thought to be consistent across these two species when abortions do occur (Schlafer & Foster, 2015).

Based on these recent reports, the absence of significant histopathologic findings in our samples may initially seem surprising. However, it is noteworthy that all of the RVFV-PCR positive placental samples that were examined in this study showed moderate to marked autolysis. Given the fact that necrosis is typically limited to the placental trophoblast layer in RVFV abortions, it seems plausible that histopathologic changes that are limited to superficial tissue layers may have been obscured by post-mortem degeneration. This finding highlights the importance of using adjunct diagnostic tests including RT-qPCR and serology to monitor for RVFV-associated abortions in livestock in areas with limited capacity for rapid sampling response and/or histopathologic examination.

**BoHV-1-associated abortions:** Two abortion events in cattle met our study case definitions for BoHV-1. In both of these cases, placentitis was observed. BoHV-1 abortion is associated with necrotic placentitis and vascular necrosis in experimental studies (Rodger *et al.*, 2007). As mentioned above, one animal, SEBI-184, that showed evidence of necrotising placentitis and vascular necrosis met both BoHV-1 and *N. caninum* case definitions and hence a co-infection cannot be ruled out. Similarly, invasive Gram-positive bacteria were seen in the placenta of the other BoHV-1 positive case suggesting that a bacterial aetiology, or co-infection, cannot be ruled out. It is also worth noting that our confidence in our study case definition for BoHV-1 is lower than for some other agents (e.g. *N. caninum*) as we relied only on evidence of seroconversion rather than direct demonstration of the pathogen. However, when taken together, our serology and histopathology results indicate that BoHV-1 is likely to be a contributing cause to these abortions.

**BVDV:** Placental pathology was not detected in the one cattle abortion event associated with BVDV in our study. This is typical of BVDV infection in cattle (Murray, 1991, Schlafer & Foster, 2015). Foetal loss and abortion associated with BVDV occurs due to foetal infection with the pathogen. However, the outcome of these foetal infection depends upon both the timing of infection and the viral subtype (e.g., cytopathic or non-cytopathic viral biotypes). For example; infection during the first trimester (up to 80 days) can result in the birth of a persistently infection calf, whereas infection between 80-150 days gestation can lead to teratogenic effects in the foetus, which in turn may result in foetal death (Lanyon *et al.*, 2014, Schlafer & Foster, 2015). Unfortunately, due to the lack of foetal tissue samples available for histopathology, we were unable to evaluate foetal pathology in this case. However, the absence of placental pathology in this BVDV-positive case is consistent with our understanding of the pathogenesis of BVDV infection in pregnant cattle.

***Coxiella burnetii*:** Histopathology and IHC findings found evidence of placentitis with positive tissue labelling by IHC in one goat abortion event (SEBI-054) that met our case definition for *C. burnetii*-associated abortion. In this case, a very high bacterial load was present in the placental tissue as indicated by the low average Ct value of 12.5 obtained with the *IS1111*a qPCR assay. Acute, diffuse, suppurative placentitis accompanied by extensive necrosis is a typical finding in *C. burnetii*-associated abortion (Schlafer & Foster, 2015) and the presence of positive IHC antibody labelling within trophoblast cells further supports the causal relationship in this case.

Positive *IS1111*a qPCR results were obtained in other six abortion events with placental material for histopathology but no other cases met the Ct cut-off based on the minimum bacterial load recommended by OIE and EFSA for diagnosing *C. burnetii*-associated abortion (see supplementary material D) (Sidi-Boumedine *et al.*, 2010, World Organisation for Animal Health (OIE), 2018). No evidence of placental pathology or substantial amounts positive tissue labelling was detected in these cases either, which supports our use of the average placental Ct cut-off ≤ 32 for attributing clinical significance to and diagnosing abortions associated with *C. burnetii* in this study. Placental tissue from one cattle abortion event (SEBI-055) did show a very small amount of granular tissue labelling, and this case also had an average placental Ct value of 32.2, very close to our diagnostic cut-off value of ≤32. This particular abortion event met study case definitions for *N. caninum* and due to the lack of placental pathology and the very low amount of IHC antibody labelling, this seems like a more likely aetiology for the observed abortion in this case. However, it remains a possibility that both pathogens were contributing to the aetiology of abortion in this case.

**Other potential aetiologies of abortion detected by histopathology:** Significant accumulations of bacteria were detected in two cattle cases (SEBI-103 and SEBI-172) that were negative by PCR or serology for all study pathogens. In both cases, moderate to high numbers of monomorphic coccobacilli were observed within the placental tissue or on the placental surface (see Table S1C). This suggests that other bacterial pathogens that were not included in our diagnostic strategy could also be making an important contribution to the problem of livestock abortion in Tanzania. Unfortunately, it was beyond the scope of this study to conduct further pathogen-specific diagnostic tests on these tissue samples, but this remains a priority question for future research in this sample set.

**S1, C; Table C:** Target pathogen serologic (ELISA) and molecular (PCR/RT-PCR) assay information, including primer/probe sequences and cycling conditions for samples from various livestock species.

| **Pathogen** | **Livestock species tested** | **Serologic test** | **PCR type** | **Primer/**  **probe name** | **Primer/probe sequence (5’ to 3’) or commercial kit** | **Source** | **Cycling conditions (temperature/time)** | | | | | |
| --- | --- | --- | --- | --- | --- | --- | --- | --- | --- | --- | --- | --- |
|  |  |  |  |  |  |  | **RT step**  **(1 cycle)** | **Initial heat inactivation**  **(1 cycle)** | **Denaturation** | **Annealing or**  **Anneal/Extension** | **Extension** | **Final extension**  **(1 cycle)** |
|  |  |  |  |  |  |  |  |  | **(35-45 cycles)** | | |  |
| *Brucella* spp. | C, G, S | Brucella COMPELISA Animal Plant Health Agency (APHA) UK | qPCR | IS7111 F | GGCCTACCGCTGCGAAT | (Matero et al., 2011) | n/a | 95 °C/  2 min | 95 °C/  5 sec | 60 °C/  30 sec | n/a | n/a |
|  |  |  |  | IS711 R | TTGCGGACAGTCACCATAATG |  |  |  |  |  |  |  |
|  |  |  |  | IS711 probe | **FAM-**AAGCCAACACCCGGC-**MGBNFQ** |  |  |  |  |  |  |  |
|  |  |  |  | IPC | Taqman® Exogenous Internal Positive Control (Applied Biosystems) |  |  |  |  |  |  |  |
|  |  |  |  | bcsp31 F | GCTCGGTTGCCAATATCAATGC | (Probert *et al.*, 2004) | n/a | 95 °C/  2 min | 95 °C/  5 sec | 60 °C/  30 sec | n/a | n/a |
|  |  |  |  | bcsp31 R | GGGTAAAGCGTCGCCAGAAG |  |  |  |  |  |  |  |
|  |  |  |  | bcsp31 probe | **FAM-**AAATCTTCCACCTTGCCCTTGCCATCA-**BHQ1** |  |  |  |  |  |  |  |
| *Chlamydia abortus* | Serology: G, S only  Molecular: C, G, S | In-house Moredun ELISA |  | CabMOMP F | GCGGCATTCAACCTCGTT | (Livingstone *et al.*, 2009) | n/a | 95 °C/  2 min | 95 °C/  5 sec | 60 °C/  5 sec | n/a | n/a |
|  |  |  |  | CabMOMP R | CCTTGAGTGATGCCTACATTGG |  |  |  |  |  |  |  |
|  |  |  |  | CabMOMP probe | **FAM** – TGTTAAAGGATCCTCCATAGCAGCTGATCAG – **BHQ1** |  |  |  |  |  |  |  |
| *Coxiella burnetti* | C, G, S | Thermofisher® PrioCHECK Ruminant Q Fever Ab Plate (ELISACOXLS5) |  | IS1111 F | CATCACATTGCCGCGTTTAC | (Roest *et al.*, 2011) | n/a | 95 °C/  2 min | 95 °C/  5 sec | 60 °C/  5 sec | n/a | n/a |
|  |  |  |  | IS1111 R | GGTTGGTCCCTCGACAACAT |  |  |  |  |  |  |  |
|  |  |  |  | IS1111 probe | **FAM** – AATCCCCAACAACACCTCCTTATTCCCAC – **BHQ1** |  |  |  |  |  |  |  |
| *Leptospira* | Serology: C only  Molecular: C, G, S | Linnodee® Bovine Leptospira Hardjo 5 Plate Solid ELISA kit (LBLK01) |  | lipL32 F | AAGCATTACCGCTTGTGGTG | (Stoddard *et al.*, 2009) | n/a | 95 °C/  2 min | 95 °C/  5 sec | 60 °C/  5 sec | n/a | n/a |
|  |  |  |  | lipL32 R | GAACTCCCATTTCAGCGATT |  |  |  |  |  |  |  |
|  |  |  |  | lipL32 probe | FAM – AAAGCCAGGACAAGCGCCG – BHQ1 |  |  |  |  |  |  |  |
| *Neospora* and  *Toxoplasma* | C, G, S | ID Screen® *Neospora caninum* Indirect ELISA    IDvet Toxoplasmosis Indirect Multi-species ELISA | 1° and 2°Nested conventional PCR | NN1 F | TCAACCTTTGAATCCCAA | (Burrells *et al.*, 2013, Buxton *et al.*, 1998, Hurtado *et al.*, 2001) | n/a | 95 °C/  5 min | 95 °C/  1 min | 55 °C/  1 min | 72 °C/  1 min | 72 °C/  5 min |
|  |  |  |  | NN1 R | CGAGCCAAGACATCCATT |  |  |  |  |  |  |  |
|  |  |  |  | NeoNP1 F | TACTACTCCCTGTGAGTTG |  |  |  |  |  |  |  |
|  |  |  |  | NeoNP1 R | TCTCTTCCCTCAAACGCT |  |  |  |  |  |  |  |
|  |  |  |  | ToxNP1 F | GTGATAGTATCGAAAGGTAT |  |  |  |  |  |  |  |
|  |  |  |  | ToxNP1 R | ACTCTCTCTCAAATGTTCCT |  |  |  |  |  |  |  |
| BTV | C, G, S | n/a | RT-PCR | Commercial | Virotype BTV pan/8 RT-PCR Kit (VT280445) Indical® | n/a | 50 °C/  20 min | 95 °C/  15 min | 95 °C/  30 sec | 57 °C/  45 sec | 68 °C/  45 sec | n/a |
| Pestiviruses | Serology: C only  Molecular: C, G, S | IDEXX® BVDV p80 Ab ELISA (P00645-5) |  | Commercial | Virotype BVDV RT-PCR Kit (VT280375) Indical® | n/a | 50 °C/  20 min | 95 °C/  15 min | 95 °C/  30 sec | 57 °C/  45 sec | 68 °C/  45 sec | n/a |
| RVF | C, G, S | ID Screen® Rift Valley Fever Competition Multispecies ELISA kit (RIFTC-10P) |  | RVS | AAAGGAACAATGGACTCTGGTCA | (Drosten *et al.*, 2002) | 45 °C/  10 min | 95 °C/  5 min | 95 °C/  5 sec | 60 °C/  30 sec | n/a | n/a |
|  |  |  |  | RVAs | CACTTCTTACTACCATGTCCTCCAAT |  |  |  |  |  |  |  |
|  |  |  |  | RVP | **FAM** -AAAGCTTTGATATCTCTCAGTGCCCCAA -**BHQ1** |  |  |  |  |  |  |  |

Where: C = cattle, G = goats, S = Sheep

**S1, D:**

**Genetic characterisation of *T. gondii* isolated from an abortion case**

**Methods:** Any samples positive for *T. gondii* by ITS1 PCR were genotyped using PCR-RFLP targeting 10 genetic markers (SAG1, SAG2 (5’-3’ SAG2 and alt.SAG2), SAG3, BTUB, GRA6, C22-8, C29-2, L358, PK1 and Apico). PCR-RFLP conditions for all markers were carried out as previously described (Hamilton *et al.*, 2015, Su *et al.*, 2010). Typing profiles were determined using RFLP banding profiles of reference strains RH (Type I), M4 (Type II variant; used for all markers except Apico where Type II strain Me49 was used instead) and NED (Type III).

**Results:** A single sheep sample was positive for *T. gondii* by PCR. Sequences was obtained for seven of 10 genetic markers. Although a complete genetic profile was not obtained, Type II alleles were identified at all seven amplified markers, indicative of a type II clonal genotype infection.

**S1; E:** Determining qPCR cut-off values for diagnosing *Coxiella burnetii* as the cause of abortion in ruminant livestock in Tanzania.

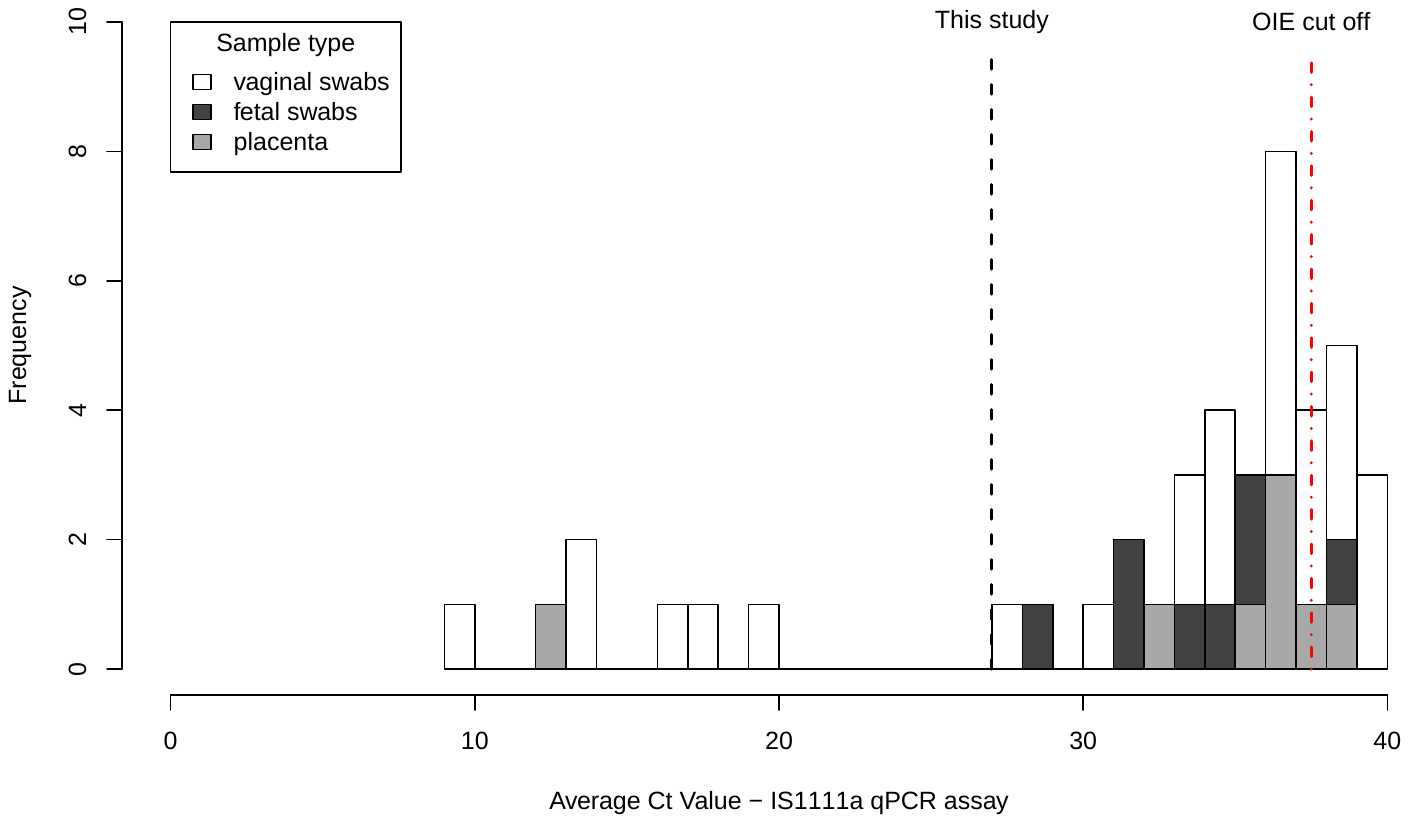

**Figure S1; E: Histogram showing frequency distribution of average Ct values obtained from the IS1111a qPCR assay for the detection of *C. burnetii* in vaginal swab, fetal swab and placental tissue samples from ruminant livestock in Tanzania**. The black dashed line indicates the cut-off Ct value (Ct value ≤ 27) selected for use in this study for attributing abortion events to *C. burnetii* infection. The red dash-dot line indicates the cut-off Ct value for vaginal swabs as stated by the OIE for attributing abortion events to *C. burnetii* infection (> 10^4^ bacteria per swab, equivalent to a Ct value = 37.5 on our platform). The equivalent cut-off value for placental tissue quoted by the OIE is off the scale (> 10^4^ bacteria per gram of placental tissue, equivalent to a Ct value = 42.7 on our platform). The histogram was plotted in R (R Core Team, 2016) using average Ct value data from all samples positive for *C. burnetii* by IS1111a qPCR assay (Klee *et al.*, 2006).

Validation of selected Ct cut-off in placental tissue samples using with histopathology and immunohistochemistry results: Histopathology and immunohistochemistry (IHC) for *C. burnetii* was performed on seven placental samples (five bovine, two caprine) that were qPCR-positive for *C. burnetii* by IS1111a qPCR testing (See Supplementary material Table S1, B). Of these, one caprine placental sample had a Ct value ≤ 27 (average Ct value of 12.5) and all other samples had Ct values > 27 (range: 32.2 - 37.5). Placentitis with positive *C. burnetii* IHC labelling was detected in the caprine placental sample with a Ct ≤ 27. No evidence of significant placental pathology or significant IHC labelling was detected in any of the other samples. These findings support our selected Ct value cut-off to support diagnosis of *C. burnetii* associated abortions within our study population.

World Organisation for Animal Health (OIE), 2018: Q Fever. In: OIE (ed.), *OIE Terrestrial Manual*
